# Mutability of mononucleotide repeats, not oxidative stress, explains the discrepancy between laboratory-accumulated mutations and the natural allele-frequency spectrum in *C. elegans*

**DOI:** 10.1101/2021.02.09.430480

**Authors:** Moein Rajaei, Ayush Shekhar Saxena, Lindsay M. Johnson, Michael C. Snyder, Timothy A. Crombie, Robyn E. Tanny, Erik C. Andersen, Joanna Joyner-Matos, Charles F. Baer

## Abstract

Important clues about natural selection can be gleaned from discrepancies between the properties of segregating genetic variants and of mutations accumulated experimentally under minimal selection, provided the mutational process is the same in the lab as in nature. The ratio of transitions to transversions (Ts/Tv) is consistently lower in *C. elegans* mutation accumulation (MA) experiments than in nature, which has been argued to be in part due to increased oxidative stress in the lab environment. Using whole-genome sequence data from a set of *C. elegans* MA lines carrying a mutation (*mev-1*) that increases the cellular titer of reactive oxygen species (ROS), leading to increased endogenous oxidative stress, we find that the base-substitution spectrum is similar between *mev-1* lines, its wild-type progenitor (N2), and another set of MA lines derived from a different wild strain (PB306). By contrast, the rate of short insertions is greater in the *mev-1* lines, consistent with studies in other organisms in which environmental stress led to an increase in the rate of insertion-deletion mutations. Further, the mutational properties of mononucleotide repeats in all strains are qualitatively different from those of non-mononucleotide sequence, both for indels and base-substitutions, and whereas the non-mononucleotide spectra are fairly similar between MA lines and wild isolates, the mononucleotide spectra are very different. The discrepancy in mutational spectra between lab MA experiments and natural variation is likely due to a consistent (but unknown) effect of the lab environment that manifests itself via different modes of mutability and/or repair at mononucleotide loci.

## Introduction

It is a fundamental principle of population genetics that the DNA sequence diversity in a population (*θ*) represents the product of mutation (*μ*) and “everything else”, where “everything else” subsumes the contributions of natural selection and random genetic drift in the composite parameter *N_e_*, the genetic effective population size: *θ*=4*N_e_μ* (Watterson 1975; Nei and Li 1979). Empirically partitioning genetic variation into the contributions of mutation and everything else requires that mutations be observed under conditions in which the effects of natural selection are minimized as much as possible. That can be done in three basic ways: by genotyping related individuals in a natural pedigree, as is now done commonly in humans (Gao et al. 2019; Halldorsson et al. 2019), by means of a “mutation accumulation” (MA) experiment, or by cataloging very rare segregating variants that have (presumably) arisen only recently and thus been minimally sieved by natural selection (Messer 2009; Carlson et al. 2018).

An MA experiment is in essence a large pedigree in which many replicate descendant lines (MA lines) are derived simultaneously from a common ancestor and allowed to evolve under conditions in which selection is minimal (Halligan and Keightley 2009). Typically, selection in an MA experiment is minimized by minimizing *N_e_*, ideally to a single individual or chromosome. Pedigree genotyping has two important advantages over MA experiments: it can be done in practically any organism, and it is fast. MA experiments are slow, labor intensive, and are only practical in organisms with fast generation times that can be maintained easily in the laboratory. However, MA experiments come with one unique advantage, which is that the phenotypic effects of mutations can be assessed under controlled experimental conditions. MA experiments have been a workhorse of evolutionary genetics for the past 60 years (Sprague et al. 1960; Mukai 1964; Liu and Zhang 2019), and much of our understanding of the mutational process has been derived from MA data (Drake et al. 1998; Halligan and Keightley 2009; Katju and Bergthorsson 2019).

The utility of MA experiments is based on a key assumption: the mutational process under laboratory conditions faithfully reflects that in nature. It has long been known from studies with microbes that all elements of the mutational process – rate, molecular spectrum, and phenotypic effects – depend on the environmental context and the genetic background; this conclusion has recently been extended to multicellular eukaryotes (Sharp and Agrawal 2012; Sharp and Agrawal 2016; Kessler et al. 2020). On the one hand the context-dependence of the mutational process is not a surprise; for example, the mutagenic effects of X-rays have been known for a century. On the other hand, it suggests that MA experiments come with their own biases (Baer 2019), just as natural selection biases the standing genetic variation.

The nematode *Caenorhabditis elegans* is an important model organism in many areas of biology, and was the first metazoan organism to have its mutational process characterized at the genomic level (Denver et al. 2000; Denver et al. 2004b). As genomic sequence data from *C. elegans* MA lines and wild isolates have accumulated, it has become apparent that the base-substitution spectrum in lab-accumulated mutations differs from that of wild isolates in a consistent way: there are more transversions in the lab than there are in nature. The ratio of transitions to transversions (Ts/Tv) in MA lines is consistently around 0.6-0.8, whereas the Ts/Tv ratio among wild isolates is around 1.1-1.2. Two early studies reported that G:C→T:A transversions are overrepresented in MA data relative to the standing nucleotide diversity (Denver et al. 2009; Denver et al. 2012); two more recent studies with more data reported that A:T→T:A transversions are also overrepresented in MA data relative to the standing nucleotide diversity (Konrad et al. 2019; Saxena et al. 2019).

The MA spectrum may differ from the standing spectrum for several reasons. First, it may be that the mutational milieu in the lab is consistently different from that in nature. Second, it may be that purifying selection against transversions in nature is stronger than against transitions, and there is some reason to think that idea is plausible (Babbitt and Cotter 2011; Guo et al. 2017). Third, it may be that the analyses of MA data and standing genetic variation come with different biases.

One key difference between the lab MA environment and the natural environment is that worms in the lab are fed *ad libitum* and kept at low population density and at a constant benign temperature, which may result in a long-term elevation of metabolic rate relative to that experienced in nature. A second key difference is that worms kept on plates in the lab experience near-atmospheric concentrations of O_2_, which is substantially greater than the optimum, as assayed by worm preference (Gray et al. 2004). Both increased metabolic rate and increased O_2_ partial pressure could potentially increase the cellular concentrations of reactive oxygen species (ROS). ROS are present in all cells as a natural byproduct of cellular metabolism, and can induce potentially mutagenic oxidative damage to DNA (Cooke et al. 2003; Bridge et al. 2014) and alterations in chromatin structure (for review, Kreuz and Fischle 2016). Accordingly, variation in cellular ROS levels has long been invoked as a potential cause of variation in mutation rate (Richter et al. 1988; Shigenaga et al. 1989; Martin and Palumbi 1993; Stoltzfus 2008). Cellular oxidative stress can vary due to differences in input (e.g., ROS levels increase under conditions of physiological stress), in the ability of cells to enzymatically convert ROS into benign products, and in repair processes (Beckman and Ames 1998; Turrens 2003; Halliwell and Gutteridge 2007; Constantini 2014).

One well-documented manifestation of oxidative damage to DNA is the oxidation of guanine to 8-oxo-7,8-dihydroguanine (8-oxodG), which if unrepaired, results in a G→T transversion (Cheng et al. 1992; Cunningham 1997; Helbock et al. 1998; Cadet et al. 2003; Evans and Cooke 2004). Accordingly, G:C →T:A transversions are often interpreted as a signature of oxidative damage to DNA (Krašovec et al. 2017; Suzuki and Kamiya 2017; Poetsch et al. 2018). While much of the early work linking ROS with 8-oxodG and mutation was conducted in somatic cells (Dollé et al. 2000; Busuttil et al. 2003), this work has been extended to sperm (e.g., Paul et al. 2011; Kim and Velando 2020). Because previous studies have found that the frequency of G:C→T:A transversions is greater in MA lines than in the standing variation, we were led to speculate that some property of the lab MA environment increases oxidative stress relative to that experienced in nature. To test that hypothesis, we employed a mutant strain of *C. elegans*, *mev-1*, that experiences elevated steady-state oxidative stress (Ishii et al. 1998; Senoo-Matsuda et al. 2001; Ishii et al. 2013). We reasoned that, if G:C→T:A transversions are in fact a signature of oxidative damage to DNA, then a strain with an endogenously increased oxidative damage should show an even greater frequency of G:C→T:A transversions than the background MA frequency.

Here we report results from an experiment in which a set of MA lines derived from a strain carrying the *mev-1(kn1)* mutation backcrossed into our canonical wild-type N2 background were propagated for ~125 generations using our standard *C. elegans* MA protocol (Joyner-Matos et al. 2011). We sequenced the genomes of 23 *mev-1* lines and compared the rate and spectrum of mutation to that of the N2 strain. As a further comparison, we report results from an additional set of 67 MA lines derived from a different wild-type strain, PB306. The laboratory MA spectrum is compared to the natural mutation spectrum inferred from “private alleles” present as homozygous variants in one and only one wild isolate (*n*=773 wild isolates).

## Results

### Mutation Rate

Mononucleotide repeats mutate differently than other sequence. We first report genome-wide rates, which are ultimately of the most evolutionary relevance; we parse mutation rates into mononucleotide and non-mononucleotide rates in the next section. The experimental design is depicted in **Figure 1**.

**Figure 1.**
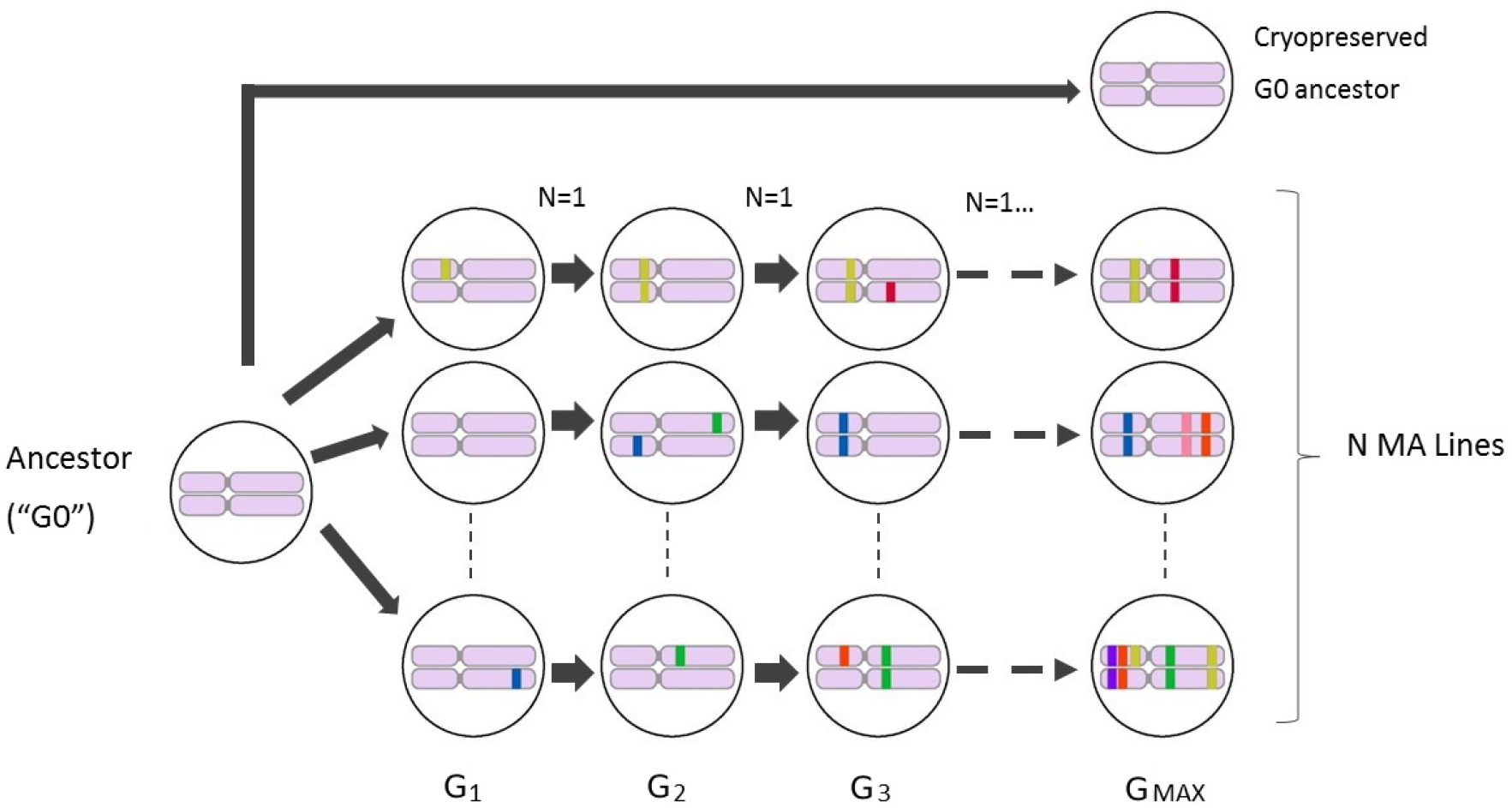
Propagation of MA lines. The common ancestor of the MA lines (G0) was thawed from a cryopreserved sample and a single immature hermaphrodite picked onto an agar plate. Lines were propagated by single-worm transfer at four-day (one generation) intervals for *t=Gmax* transfers. Lines were initially genetically homogeneous (pink chromosome pairs); colored bars represent new mutations, which are fixed in a line with expectation *u* =(1-*s*)/(2-*s*), where *s* is the selection coefficient (Keightley and Caballero 1997). See Methods and **Supplemental Table S1** for details of the MA experiments.

#### (i) Genome-wide mutation rate

Summary statistics on nuclear mutation rates are presented in **Table 1** and results of statistical tests in **Supplemental Table S2**. Results for individual MA lines are given in **Supplemental Table S3**. A complete list of mutations and their genomic context is given in **Supplemental Table S4 (nuclear loci)** and **Supplemental Table S5 (mtDNA loci)**. Raw sequence data are archived in the NCBI Short Read Archive, project numbers PRJNA429972 (32 N2 MA lines) and PRJNA665851 (all other MA lines).

**Table 1.** Mutation rates. (*μ*) x 10^9^/site/generation (SEM). *U_GENOME_* is the haploid genome-wide mutation rate per-generation.

|  | *Mev-1* (*n*=23 MA lines) |  |  | N2 (*n*=68 MA lines) |  |  | PB306 (*n*=67 MA lines) |  |  |
| --- | --- | --- | --- | --- | --- | --- | --- | --- | --- |
|  | *non-mono* | *mono* | *Total* | *non-mono* | *mono* | *Total* | *non-mono* | *mono* | *Total* |
| $\mu_{GC \rightarrow AT}$ | 1.45 (0.12) | 1.42 (0.36) | 1.45 (0.10) | 1.83 (0.06) | 1.6 (0.24) | 1.81 (0.06) | 1.81 (0.05) | 1.91 (0.22) | 1.82 (0.05) |
| $\mu_{GC \rightarrow TA}$ | 0.81 (0.12) | 1.29 (0.36) | 0.85 (0.11) | 0.80 (0.04) | 1.13 (0.18) | 0.82 (0.04) | 0.94 (0.04) | 1.68 (0.21) | 1.00 (0.05) |
| $\mu_{GC \rightarrow CG}$ | 0.41 (0.06) | 0.65 (0.26) | 0.43 (0.06) | 0.46 (0.04) | 0.42 (0.09) | 0.46 (0.03) | 0.43 (0.03) | 0.42 (0.09) | 0.43 (0.03) |
| $\mu_{AT \rightarrow GC}$ | 0.39 (0.05) | 0.33 (0.12) | 0.38 (0.04) | 0.47 (0.03) | 0.40 (0.05) | 0.46 (0.02) | 0.45 (0.03) | 0.34 (0.04) | 0.43 (0.02) |
| $\mu_{AT \rightarrow CG}$ | 0.20 (0.03) | 0.32 (0.10) | 0.22 (0.03) | 0.29 (0.02) | 0.39 (0.05) | 0.31 (0.02) | 0.25 (0.02) | 0.35 (0.04) | 0.27 (0.02) |
| $\mu_{AT \rightarrow TA}$ | 0.35 (0.05) | 3.04 (0.29) | 0.83 (0.06) | 0.44 (0.03) | 3.34 (0.16) | 0.95 (0.04) | 0.40 (0.02) | 2.70 (0.14) | 0.81 (0.03) |
| $\mu_{BS}$ | 1.60 (0.07) | 3.62 (0.26) | 1.89 (0.07) | 1.91 (0.05) | 3.93 (0.15) | 2.20 (0.05) | 1.89 (0.05) | 3.52 (0.16) | 2.13 (0.05) |
| $\mu_{INS}$ | 0.15 (0.03) | 1.32 (0.19) | 0.32 (0.04) | 0.07 (0.01) | 1.07 (0.06) | 0.21 (0.01) | 0.10 (0.01) | 1.19 (0.09) | 0.26 (0.02) |
| $\mu_{DEL}$ | 0.15 (0.02) | 1.76 (0.23) | 0.38 (0.03) | 0.19 (0.01) | 1.43 (0.09) | 0.36 (0.02) | 0.26 (0.02) | 1.76 (0.12) | 0.47 (0.03) |
| $\mu_{INDEL}$ | 0.30 (0.05) | 3.09 (0.22) | 0.70 (0.04) | 0.25 (0.01) | 2.50 (0.11) | 0.58 (0.02) | 0.36 (0.02) | 2.95 (0.17) | 0.73 (0.03) |
| $\mu_{TOTAL}$ | 1.90 (0.10) | 6.70 (0.41) | 2.59 (0.08) | 2.17 (0.05) | 6.43 (0.20) | 2.78 (0.06) | 2.25 (0.06) | 6.47 (0.22) | 2.85 (0.07) |
| $U_{GENOME}$ | 0.16 | 0.10 | 0.25 | 0.18 | 0.09 | 0.27 | 0.19 | 0.09 | 0.27 |

To test the hypothesis that the overall base-substitution mutation rate *μ_BS_* differs between strains, we used a general linear model (GLM); details are presented in the Methods. The hypothesis that the rate of G:C→T:A transversions is greater in *mev-1* than in N2 is a prior one-tailed hypothesis test. The GLM revealed significant variation in *μ_BS_* among the three strains, as well as a significant interaction between strain and mutation type. Contrary to our expectation, *μ_BS_* summed over all six mutation types is significantly less in the *mev-1* lines than in N2, and marginally lower than in PB306. In contrast, *μ_BS_* does not differ between N2 and PB306. The G:C→T:A transversion rate differs among strains (**Figure 2A**), but not in the predicted way: *μ_GC→TA_* is indistinguishable between *mev-1* and N2, and lower than in PB306.

**Figure 2.**
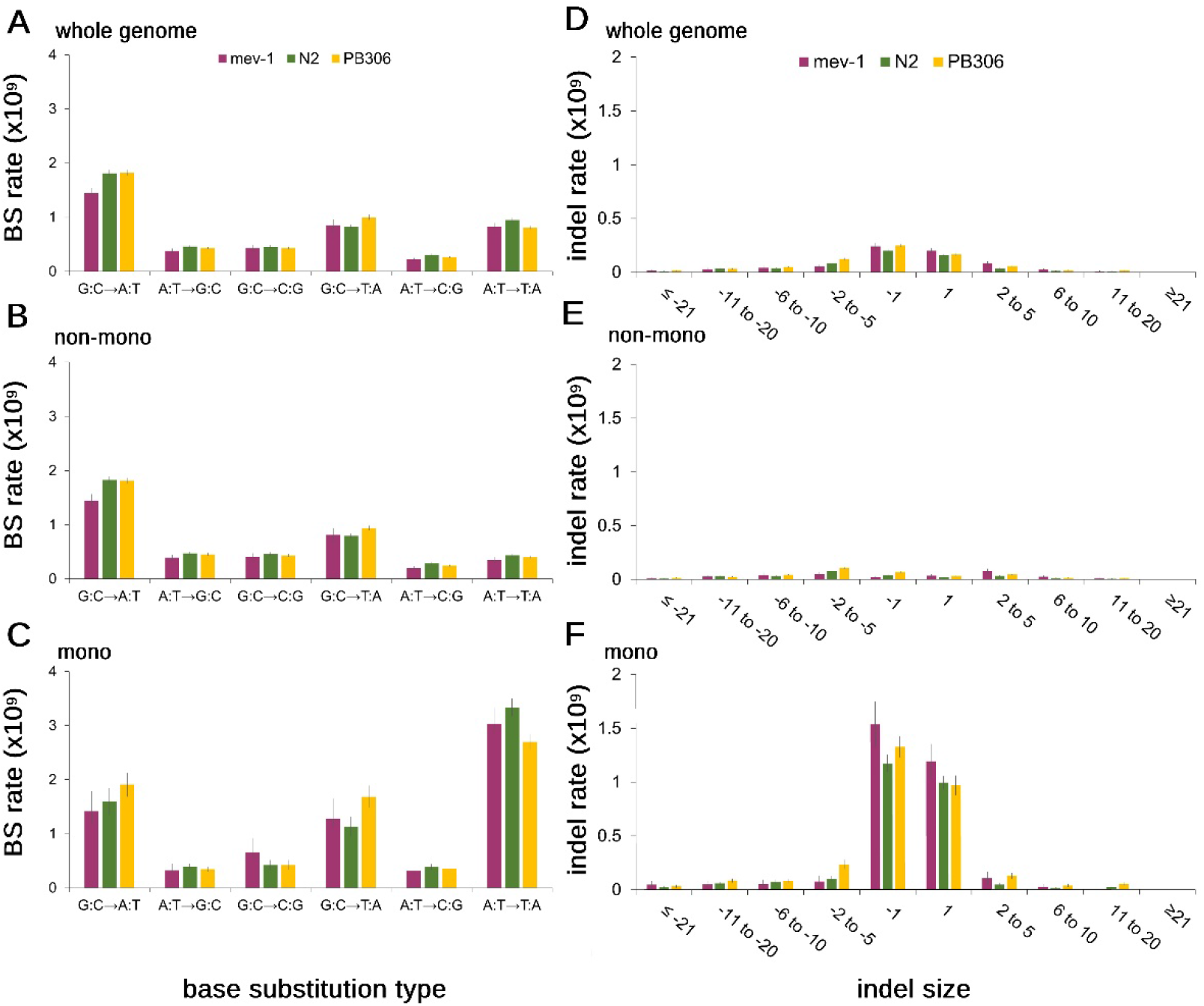
Mutation rates. Left (**A-C**), type-specific base-substitution rates, by strain. Right (**D-F**), size-specific indel rates, by strain. Top row (**A,D**), genome-wide mutation rates; middle row (**B,E**), rates at non-mononucleotide repeat sequence; bottom row (**C,F**), mononucleotide repeat sequence. All rates are scaled as 10^-9^ per base per generation; note the difference in y-axis scale between panels (**A-C**, left) and panels (**D-F**, right). Error bars represent 1 SEM.

The rate of A:T→T:A transversions is marginally different between the three strains (**Figure 2A**), with N2 having a slightly greater rate than *mev-1* and PB306.

The insertion and deletion rates both vary among strains, in different ways (**Figure 2D; Table 1**). The PB306 deletion rate is significantly greater than that of *mev-1* and N2. The *mev-1* insertion rate is significantly greater than that of N2 and greater than that of PB306, although not significantly. The PB306 insertion rate is marginally greater than that of N2.

The mechanisms of mutagenesis and repair are not necessarily uncorrelated, and the large number of mutations (>9,000) provides an opportunity to investigate the covariance structure of the different types of mutation. We first investigated the covariance structure of the six types of base-substitution. We initially fit a model with a single variance component, i.e., a uniform diagonal element in the covariance matrix and all off-diagonal elements constrained to zero. That model was compared to a model in which a separate variance (diagonal element) was estimated for each of the six traits (“banded main diagonal”) and the off-diagonal elements constrained to zero. That model was in turn compared to a model in which all elements are unconstrained. The unconstrained model had the smallest AICc (ΔAICc=16.2 less than the banded main diagonal), and provided a significantly better fit to the data (LRT, −2ΔlnL=47.1, df=15, P<0.0001), although the individual correlations were small (all |*r*|<0.3; **Supplemental Table S6**). A model with the covariance matrix estimated separately for each strain did not fit as well (ΔAICc=12.4 greater than the model with a single covariance matrix).

We next investigated the covariance structure of indels and base-substitutions together, combining all six base-substitutions into one category with insertions and deletions included separately. The best model is the most parameter-rich, with the unstructured covariance matrix estimated separately for each strain (ΔAICc=12.3 less than the model with a single unstructured covariance matrix; LRT, −2ΔlnL=37.6, df=12, P<0.0002). In all three strains, correlations between base-substitution and deletion rates are moderately positive (*r*=0.2-0.3), and base-substitution and insertion rates are uncorrelated (r≈0). The correlation between base-substitution rate and insertion rate is negative in *mev-1* and N2, whereas it is moderately positive in PB306 (*r*≈0.3; **Supplemental Table S7**). We emphasize that, for both of these covariance analyses, we cannot reject the hypothesis that any specific off-diagonal element is different from zero. Rather, the results simply mean that models with some non-zero off diagonal elements fit the data better than a model in which all off-diagonal elements are constrained to zero, and we interpret the point estimates of the individual pairwise correlations heuristically.

Inspection of the residuals of the GLM reveals two MA lines that are obvious outliers. Line 572 (N2) has anomalously high base-substitution and deletion rates, although its insertion rate is slightly below the N2 average. Line 481 (PB306) has anomalously high insertion and deletion rates, although its base-substitution rate is only slightly greater than the PB306 average (**Supplemental Table S3**). We searched the list of mutations in those two lines for candidates that could potentially increase the mutation rate (see **Extended Methods, Supplemental Appendix A1.4**). It turns out there are obvious candidates in both lines (**Supplemental Table S4**). Line 572 has a T→G transversion upstream of the *atl-1* gene; *atl-1* is involved in homologous recombination and the DNA damage checkpoint. Line 471 has a 1-bp deletion downstream of the *xpc-1* gene; *xpc-1* is involved in nucleotide excision repair.

The mitochondrion is the site of the electron transport chain subunit II, and therefore the source location of the excess free radicals produced in *mev-1* worms (Senoo-Matsuda et al. 2001). However, the mtDNA mutation rate *μ_Mt_* is not increased in *mev-1*, and is very close to that of N2 (*mev-1 μ_Mt_* =5.62+2.62×10^-8^/gen; N2 *μ_Mt_* =6.05+1.36×10^−8^/gen; randomization test p>0.86). The point estimate of the mtDNA mutation rate is ~50% greater in PB306 (*μ_Mt_ =*8.84±2.15×10^-8^/gen), but the difference between PB306 and N2 is not significantly different from zero (randomization test p>0.47). A list of mtDNA mutations and their heteroplasmic frequencies is given in **Supplemental Table S5**. Details of the calculation of *μ_Mt_* and of the randomization test are given in the Methods.

#### (ii) Mononucleotide repeats

Mononucleotide repeats are well-known to incur indel mutations at a much greater rate than non-repeat sequence (Denver et al. 2004a), and we previously found that mononucleotide repeats in the N2 genome of eight or more consecutive bases also experience an elevated rate of base-substitution mutations (Saxena et al. 2019). Here we find the same result when mononucleotides are defined by the less stringent criterion of ≥5 consecutive bases. Mononucleotide repeats of ≥5 bases comprise ~8.5% of the N2 genome, of which ~80% are A:T repeats (see **Extended Methods, Supplemental Appendix A1.8**). Pooled over strains and mutation types, the mononucleotide base-substitution rate is nearly twice that of non-mononucleotide sequence (**Table 1**). However, the increased base-substitution mutation rate in mononucleotide repeats is not uniform; the rate of transversions from either a G:C or an A:T base pair to an A:T base pair is greater in mononucleotide repeats, whereas rates of the other four types of base-substitution do not differ between sequence types (**Figure 2B,C**). In particular, the rate of A:T→T:A transversions is approximately seven-fold greater in mononucleotide repeats than in non-repeat sequence.

As expected, the rate of +/− 1 bp indel mutations is much greater in mononucleotide repeats than in non-repeat sequence; averaged over strains, 1 bp deletions occur about 26 times more frequently and 1 bp insertions about 39 times more frequently in mononucleotides than in non-mononucleotide sequence. In contrast, the rate of both deletions and insertions longer than 1 bp is only about twice as great in mononucleotides as in non-mononucleotide sequence (**Figure 2D-F; Table 1**). Summed over sequence types and mutation types, the rate of 1 bp indels is about 25% greater in *mev-1* than in N2, whereas the rates in N2 and PB306 are similar.

### Local sequence context

A previous analysis of the factors potentially affecting base-substitution mutability in N2 revealed a predominant role for the local (three-nucleotide) sequence context (Saxena et al. 2019). In that study, motifs with C or G in the mutant (3’) site were more mutable on average than motifs with A or T in the mutant site, with one exception: the triplet 5’-ttA-3’ was the most mutable of any of the 64 motifs. Subsequent MA studies with the N2 strain of *C. elegans* have found a similarly high mutability of the 5’-ttA-3’ motif (Konrad et al. 2019; Volkova et al. 2020).

The 64 nucleotide triplets are depicted in **Figure 3**, oriented 5’→3’ with the mutant site in the 3’ position. There are not sufficiently many mutations to permit a robust statistical comparison between strains of the full set of 64 triplets, but we have a prior hypothesis with respect to the relative mutability of the 5’-ttA-3’ triplet. 5’-ttA-3’ is also the most mutable motif in the other two strains, although the size of the anomaly is not as extreme in the other strains as in N2. The 5’-ttA-3’ mutation rate differs among strains (N2, *μ_ttA_*=8.6×10^-9^/gen; *mev-1*, *μ_ttA_*=6.6 ×10^-9^/gen; PB306, *μ_ttA_*=6.4×10^-9^/gen); the difference between PB306 and N2 is statistically significant. The correlation of the 64 motif-specific mutation rates between strains is large and positive (>0.7), but significantly less than 1 in all three cases (**Supplemental Table S8**).

**Figure 3.**
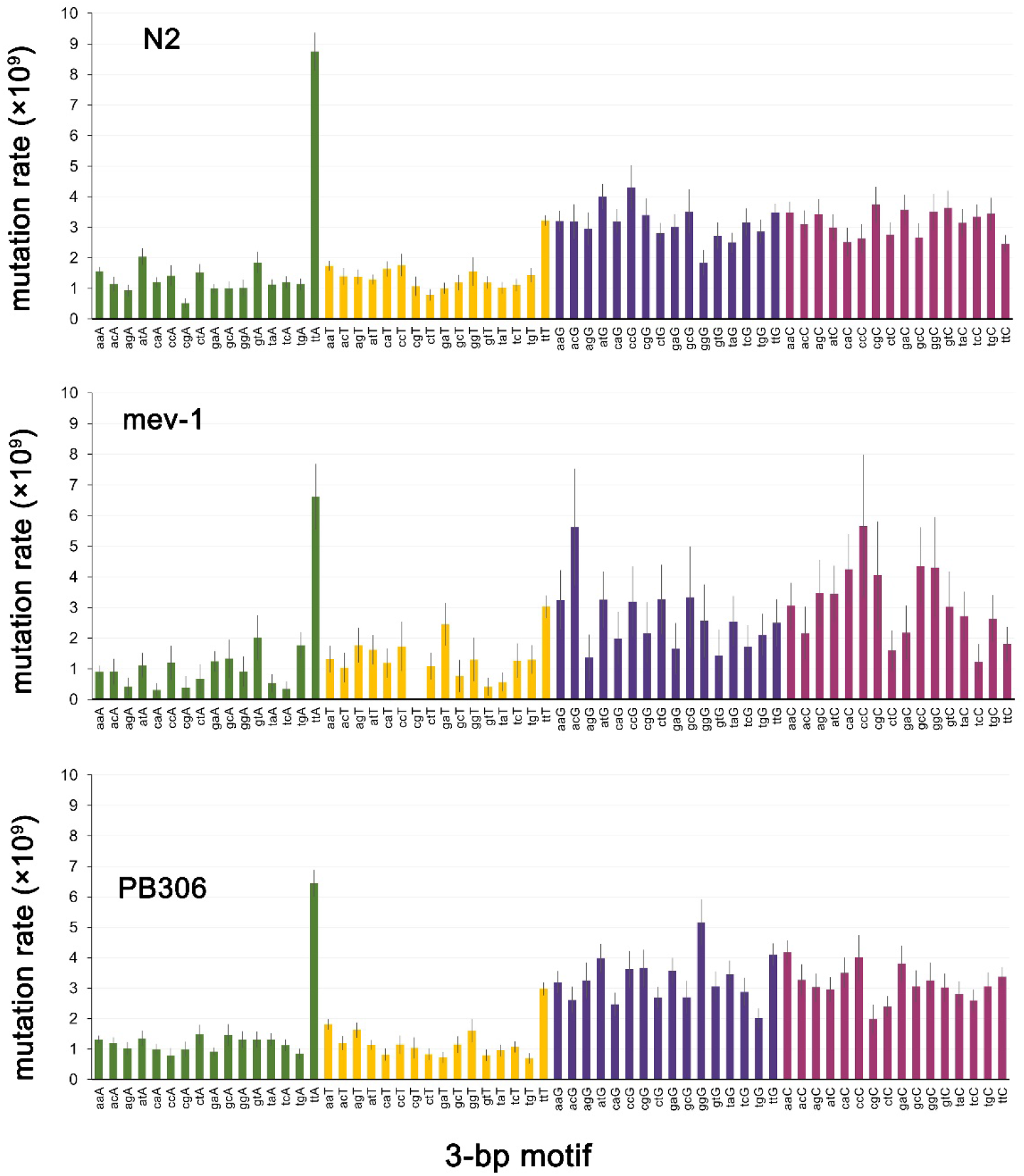
64 3-bp motif base-substitution mutation rates. Motifs are arranged 5’-xyZ-3’, with the mutant base Z in the 3’ position. Error bars show one SEM.

Some fraction of 5’-ttA-3’ triplets occur at the 3’ end of poly-T repeats, suggesting the possibility that the atypically high mutability of that motif results from the association of the motif with mononucleotide repeats. Averaged over the three strains, *μ_ttA_* when the motif is adjacent to a poly-T repeat is ~27-fold greater than when the same motif is not adjacent to a poly-T repeat (43.0 + 2.3 ×10^-9^/generation vs. 1.6+ 0.2 ×10^-9^/generation). When not in the context of a mononucleotide repeat, the 5’-ttA-3’ motif mutates at essentially the same rate as any A:T base-pair (**Figure 3**).

Details of the local sequence context analysis are presented in the **Extended Methods, Supplemental Appendix A1.9**.

### Mutation Spectrum

The mutation spectrum, defined as the frequency distribution of mutations of given types, is not the same as the distribution of type-specific mutation rates, although the two are obviously related. In our MA data, pooling over strains and the six base-substitution types, the correlation between the type-specific base-substitution mutation rate (*μ_i_*) and the proportion of mutations of that type (*p_i_*), *r_μ,p_*= 0.85. The utility of knowing the mutation spectrum in MA lines is that it provides a benchmark for comparison to wild isolates, for which the mutation rate cannot be known.

Summed over all MA lines within strains, the six-category base-substitution mutation spectrum of *mev-1* (*n*=508 mutations) does not differ from those of either N2 (*n*=3,434 mutations; 2×6 Monte Carlo Fisher’s Exact Test, 10^6^ iterations, P>0.50) or PB306 (*n*=3,111 mutations; MC FET, P>0.45), whereas the spectrum of N2 differs significantly from that of PB306 (MC FET, P<0.001; **Figure 4A**). The differences between the base-substitution spectra of N2 and PB306 are not great, but the large number of mutations enables us to detect significant differences on the order of a few percent.

**Figure 4.**
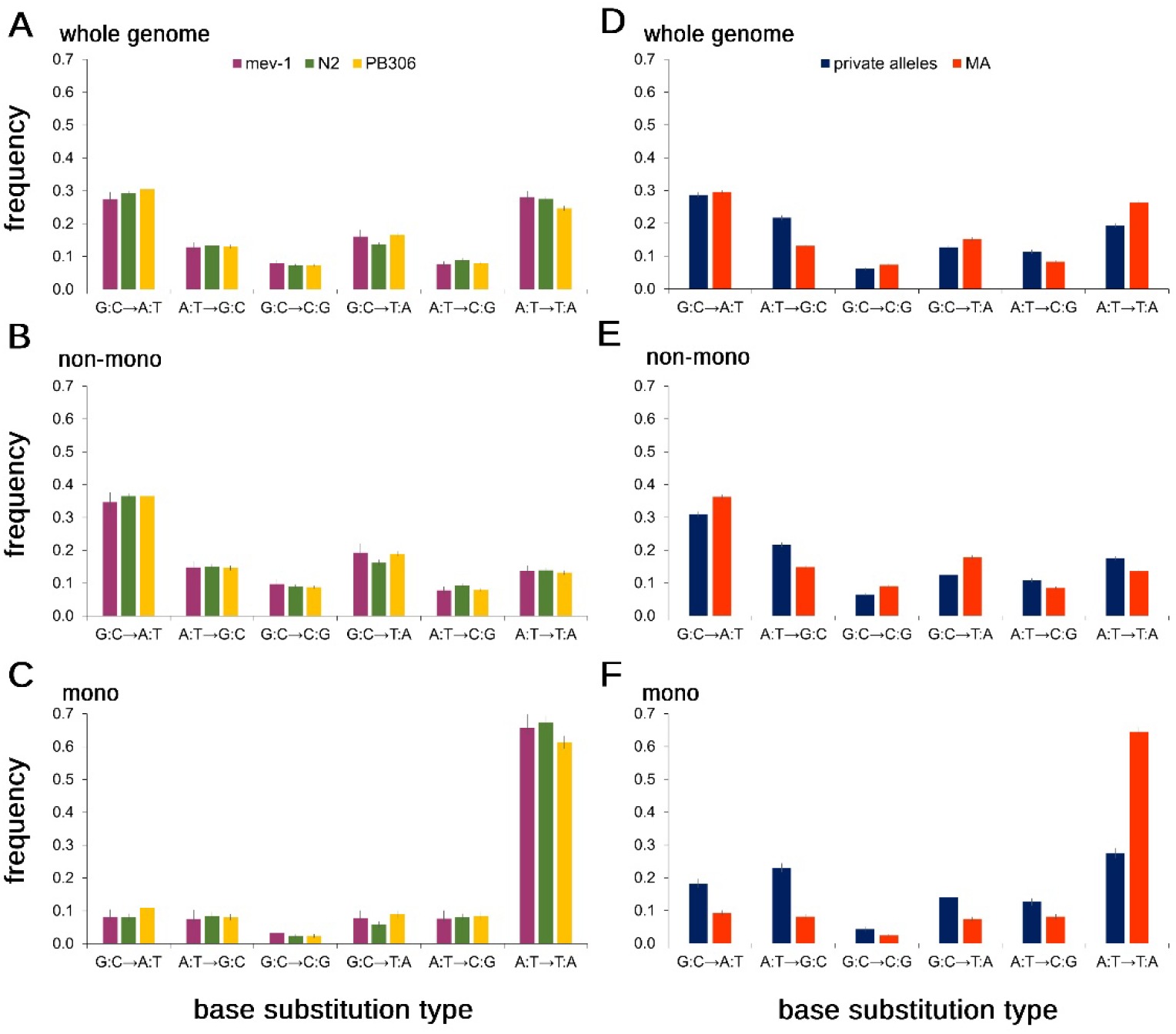
Base-substitution spectra. Left panel (**A-C**), MA lines; Right panel (**D-F**), wild isolate private alleles (blue) and MA means (red). Top panels (**A,D**), whole-genome; middle **panels (B,E**), non-mononucleotide sequence; bottom panels (**C,F**), mononucleotide sequence. Error bars show one SEM.

To compare the indel spectra between strains, we first investigated the overall mutational bias, quantified as the proportion of deletions among indel mutations. The indel bias differed significantly among strains (3×2 Fisher’s exact test, P<0.04). Breaking down the bias into pairwise comparisons between strains, *mev-1* (55% deletion) had a significantly lower proportion of deletions than N2 (63% deletion, Fisher’s Exact Test P<0.04) and PB306 (65% deletion, FET P<0.02); the bias did not differ significantly between N2 and PB306.

To characterize the indel spectra with finer resolution, we next assigned insertions and deletions separately to four bins of size 1, 2-5, 6-10 and 11-20. The bin sizes are obviously arbitrary, but the distribution provides cell counts of five or more observations for all but the largest insertion bin in the *mev-1* strain. The genome-wide spectrum (**Figure 5A**) did not differ between *mev-1* and N2 (FET, p>0.08) nor between N2 and PB306 (FET, p>0.29).

**Figure 5.**
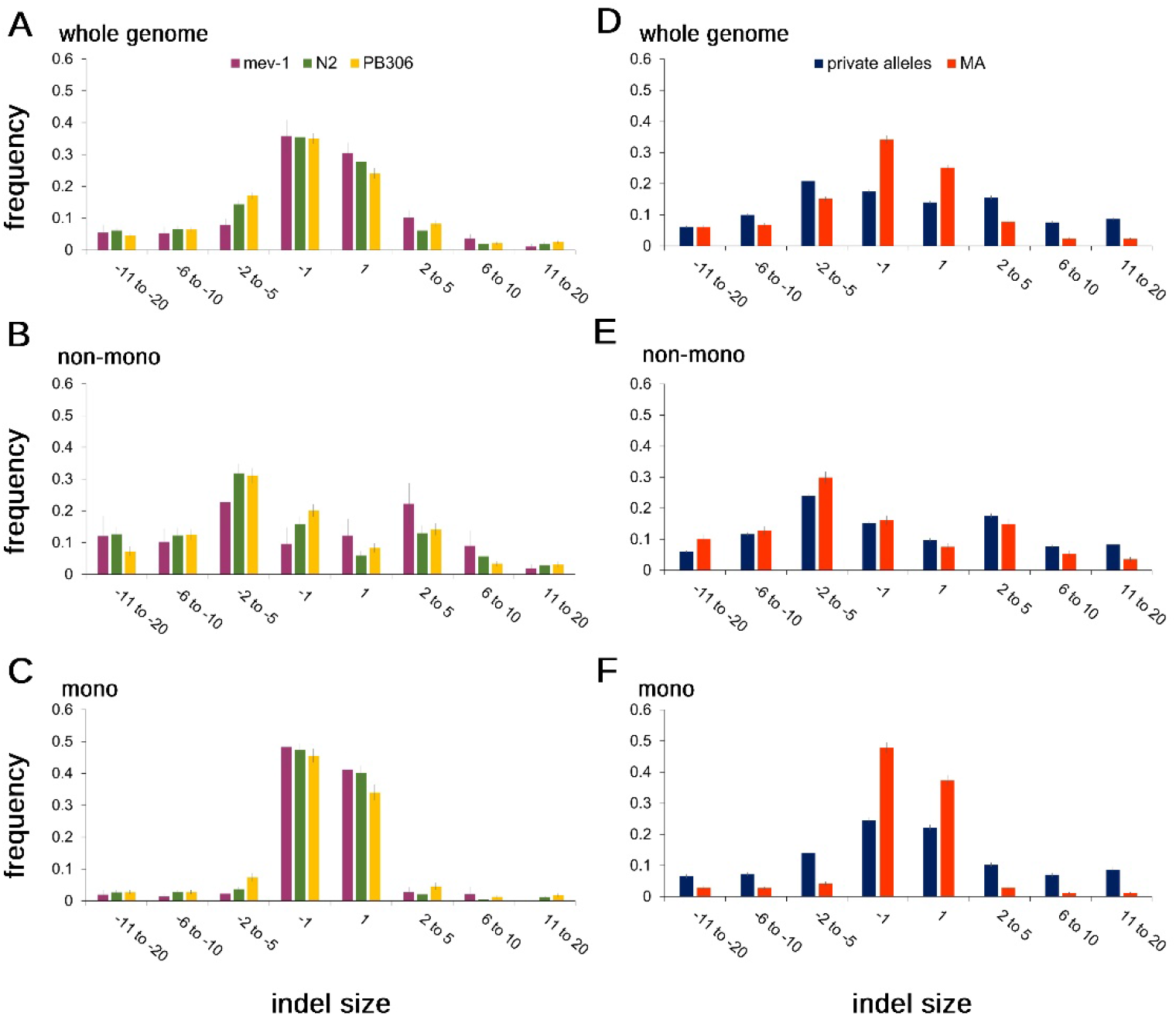
Indel spectra. Left panel (**A-C**), MA lines; Right panel (**D-F**), wild isolate private alleles (blue) and MA means (red). Top panels (**A,D**), whole-genome; middle panels (**B,E**), non-mononucleotide sequence; bottom panels (**C,F**), mononucleotide sequence. Error bars show one SEM.

Breaking the genomic context into mononucleotide repeats and non-mononucleotides, the non-mononucleotide spectra (**Figure 5B**) were marginally different between *mev-1* and N2 (FET, p<0.03), but not between N2 and PB306 (FET, p>0.11). The mononucleotide spectra (**Figure 5C**) were very similar between *mev-1* and N2 (FET, p>0.82), but significantly different between N2 and PB306 (FET, p<0.01). Averaged over N2 and PB306, the deletion bias is stronger in non-mononucleotide sequence (72.5% deletion) vs. mononucleotides (58% deletion).

### Comparison of the MA spectrum to the natural private allele spectrum

#### (i) Base-substitutions

Variant alleles present in a single wild isolate (“private alleles”) presumably arose as new mutations in the recent past, and have been minimally scrutinized by selection. As expected from previous studies, the genome-wide base-substitution spectrum of unique variants is qualitatively different from those of the MA lines (**Figure 4B**). Because of the large sample sizes, even small differences in proportions between groups are highly statistically significant (FET, P≪0.0001 in all cases), so we interpret the differences heuristically. We first consider the genome-wide spectrum, then parse the genome into mononucleotide repeats and non-mononucleotide repeats. The wild isolate spectra are compared to the average of the N2 and PB306 MA spectra (which are themselves significantly different). The full CeNDR wild-isolate dataset includes 773 unique genotypes, which collectively harbor >450,000 private single-nucleotide variants. Of those 773 wild isolates, we arbitrarily restricted our analysis to 447 isolates which carry 100 or fewer private alleles (*n*=11,667 private alleles), which represent about 400 generations of mutation accumulation. A list of the wild isolates included and the set of private alleles for each isolate are given in **Supplemental Table S9**.

A:T→T:A transversions occur more frequently in the MA lines than in the wild isolates, and A:T→G:C transitions occur less frequently in the MA lines. G:C→T:A transversions, the original focus of this study, occur only slightly more frequently in the MA lines than among the rare variants. The wild isolate private allele spectrum is closer to that of PB306 (and *mev-1*) than it is to that of N2, but the MA spectra are clearly more similar to each other than either is to the wild isolate spectrum.

Comparison of the base-substitution spectra of mononucleotide repeat sequences to the spectra of non-mononucleotide sequences is revealing (**Figure 4B**). For non-mononucleotide sequence (~90% of the genome), the private allele spectrum of wild isolates is similar to that of the MA lines, although the proportion of private alleles mutating from a C:G base-pair is less than the MA frequency in all three cases. In contrast, the mononucleotide spectra are very different. The proportion of A:T→T:A transversions in the wild isolates is less than half that expected from the MA proportion (28% vs. 65%), and there are many fewer A:T→G:C transitions in the MA lines than in the private alleles (8% vs. 25%)

If the strength of purifying selection varies among the different types of mutations, we expect the spectrum of common alleles will differ from that of rare variants. We compared the genome-wide allele frequency spectrum between private alleles and common(er) alleles, defined as variants present in more than one isolate but with an upper bound of 10% on the minor allele frequency. We chose 10% as the upper-bound on the minor allele frequency to minimize the chance of misidentifying an ancestral allele as the recent mutant. The spectra of rare and common variants are similar (**Supplemental Figures S1, S2**), reinforcing the notion that the difference in spectra between standing variants and mutations accumulated in laboratory MA experiments results from a consistent difference between the mutational milieu in the lab and in nature, rather than from the action of differential purifying selection.

As noted, the MA spectrum differs slightly but significantly between N2 and PB306. N2 is long-adapted to the lab environment, and differs from wild-type *C. elegans* in many respects (Sterken et al. 2015), of which the mutation spectrum is but one. Private alleles in the wild isolates provide a much broader sample of the mutational process in the species. Parametric bootstrap simulations (see Methods for the details) show that there is more variation in the spectrum of private alleles among wild isolates than expected from a uniform base-substitution mutational process (**Supplemental Figure S3**), both for non-mononucleotide sequence and mononucleotide runs, as well as genome-wide. This finding demonstrates that, unsurprisingly, the *C. elegans* mutational process varies among strains in nature as well as in the lab, although the causes of the variation in nature are uncertain.

#### (ii) Indels

The frequencies of false positives and false negatives are qualitatively greater for indels of >20 bp than for smaller indels (see next section), so we restrict comparison of indel spectra of MA lines and wild isolates to indels of <21 bp. To compare the indel spectra of MA lines and natural private alleles we applied additional filters on the MA lines and used the full set of 773 wild isolates (**Extended Methods, Supplemental Appendix A1.11**).

As noted previously, the MA lines have a strong genome-wide deletion bias (~62% deletion); the deletion bias in the wild isolates is much weaker (54% deletion). Inspection of the genome-wide spectrum (**Figure 5D**) reveals that +/− 1 bp indels are much a much greater fraction of indels in the MA lines than in the wild isolates. Breaking the genome into mononucleotide and non-mononucleotide components, it is apparent that the non-mononucleotide spectra are roughly congruent between MA lines and wild isolates (**Figure 5E**). In contrast, the mononucleotide spectra (**Figure 5F**) are different between MA and wild isolates, with the MA lines having a greater fraction of +/− 1 bp indels. Evidently, the genome-wide discrepancy between the MA indel spectrum and the private allele spectrum of wild isolates is largely due to the different mutational properties of mononucleotide repeats in lab populations and in nature.

### False positives and false negatives

For an estimate of mutation rate to be believable, it is necessary to have credible estimates of the frequency of false positives (FP, apparent variants called as new mutations that are not new mutations) and false negatives (FN, mutated sites in the genome that are not called as new mutations). We designate alleles that are homozygous in the *C. elegans* reference genome (N2) as “0” variant alleles are designated as “1”. Some sites in the common ancestors of our MA lines differ from the reference allele, i.e., the MA ancestor genotype is 1/1 (1,439 sites in the N2 ancestor, 1,642 in the *mev-1* ancestor, 155,614 in the PB306 ancestor).

False positives were assessed in two ways. First, we counted sites that were scored 1/1 in our MA ancestor and 0/0 in the reference genome. These are presumed to be new mutations fixed subsequent to the divergence of the MA ancestor and the reference strain from their common ancestor (alternatively, they could be errors in the reference genome). Then, we scored those sites in each MA line in the set. If a site was scored 1/1 in the MA ancestor but scored 0/0 in all MA lines, we inferred a false positive in the MA ancestor. By that criterion, the false discovery rate (FDR=FP/[FP+TP]) in the N2 MA lines for base-substitution mutations is 0.19% and in PB306 it is 0.0033%. FDRs for indels in the two strains are 0 and 0.28%, respectively.

Second, we employed an independent set of low-coverage sequence data from a set of 192 recombinant inbred advanced intercross lines (RIAILs) generated from a cross between two N2 strain MA lines, 530 and 563. The details of the construction and sequencing of the RIAILs and data analysis are presented in **Supplemental Appendix A2**; RIAIL genotypes are given in **Supplemental Table S10**. Each parental line carries its own set of putative mutations, which are expected to segregate in the RIAILs with an average frequency of 50%. A variant called as a mutant in a parental line (1/1) that does not segregate in the RIAILs is designated as a false positive in the parent. The false discovery rate for detection of SNPs and indels is 2.1% and 2.9%, respectively. This result is consistent with a previous estimate of the FDR from N2 strain MA lines based on Sanger sequencing of PCR products, which gave an upper 95% confidence limit on the FDR of 2.5% (Saxena et al. 2019).

True mutations may not be identified as such for two reasons. First, the mutant site may not be covered by the sequencing. We refer to this situation as “failure to recall” a mutation. Or, a true mutation (i.e., 0/0 in the ancestor and 1/1 in the MA line carrying the mutation) may be incorrectly called as “not 1/1”, i.e., either as 0/0 or as a heterozygote, 0/1. These are false negatives. False negatives will cause the mutation rate to be underestimated. Failure to recall mutations will only cause the mutation rate to be misestimated (under or over) if the mutation rate at sites not covered in the sequencing differs from that of the sites covered.

We employed a simulation approach to assess recall rate, the details of which are given in the **Extended Methods, Supplemental Appendix A1.10**. “Dummy" mutations were introduced into the reference genome at random, and the MA data analyzed using our standard variant-calling pipeline but with the simulated genome as the reference. Sites with dummy mutations are scored as 1/1 in the MA ancestor and all MA lines. If a dummy mutation at a site was not called 1/1 in an MA line, for any reason, it was classified as a failure to recall. The genome-wide base-substitution failure-to-recall rate is 6.81% in N2, 6.92% in *mev-1*, and 9.18% in PB306. The small indel (≤50bp) failure-to-recall rates are 9.34% in N2, 9.78% in *mev-1*, and 12.35% in PB306.

Dummy mutations that are called as “not 1/1” (i.e., 0/0 or 0/1) are false negatives *sensu stricto*. The genome-wide base-substitution false negative rate (FNR) is 0.12% in N2, 0.17% in *mev-1*, and 0.91% in PB306. The small indel (≤50bp) FNRs are 5.15% in N2, 5.59% in *mev-1*, and 5.84% in PB306.

## Discussion

Four key findings emerge. First, the base-substitution mutational process of *mev-1* is similar to that of N2, from which it was derived. Therefore, any discrepancy between the base-substitution spectrum of N2 and those of wild isolates is unlikely to be due to differences resulting from endogenous oxidative stress manifested in the lab environment. Moreover, the overall base-substitution rate of *mev-1* is significantly less (~14%) than that of N2, contrary to the prediction that elevated levels of ROS increase the base-substitution mutation rate. In contrast, the indel rate, and especially the insertion rate, is greater in the *mev-1* lines than in N2 (~50% greater). We have previously shown that the dinucleotide repeat indel rate increases in N2 MA lines propagated under conditions of heat stress (Matsuba et al. 2013), and a similar increase in the indel rate – but not the base-substitution rate - has been shown in *Arabidopsis thaliana* MA lines cultured under conditions of heat stress (Belfield et al. 2020). *Drosophila melanogaster* MA lines initiated from genomes carrying deleterious large-effect mutations experience higher rates of short deletions than wild-type MA lines, thought to be due to preferential deployment of different mechanisms of double-strand break repair (Sharp and Agrawal 2016). The increased insertion rate in the *mev-1* lines suggests that elevated levels of endogenous oxidative damage may in fact influence the mutational process, but not in the way that we predicted.

A possible explanation for the lower base-substitution rate in *mev-1* relative to N2 is that the N2 lines underwent approximately twice as many generations of MA (Gmax=250 generations) as did *mev-1* (Gmax=125 generations). We previously found that “second order” N2 MA lines initiated from a subset of the N2 lines included in this study evolved a significantly greater (~10%) mutation rate over a subsequent ~150 generations of MA. The results reported here are consistent with the N2 (and PB306) lines evolving an increased base-substitution mutation rate over time, although the evidence is circumstantial.

The second key result emerges from the comparison of the mutational properties of the lab-adapted N2 strain and the wild isolate PB306. The overall base-substitution mutation rates are very similar between the two strains (<5% different), although some of the type-specific mutation rates are marginally different (e.g., the genome-wide G:C→A:T transition rate is ~20% greater in PB306, and the A:T→T:A transversion rate is ~20% greater in N2). The type-specific differences between strains are not large – by way of contrast, two sets of *D. melanogaster* MA lines derived from different starting genotypes had a five-fold difference in G:C→A:T transition rates (Schrider et al. 2013). PB306 has a similarly greater indel rate than N2 (~25%), although the direction and magnitude of the bias are nearly identical (65% and 63% deletion, respectively).

The differences in mutational properties between N2 and PB306 surely have a genetic basis. The two strains’ genomes are about 1.5% different (~150K pairwise differences). With only two strains, we cannot say whether the difference is related to the domestication of N2, although it is interesting that the mutational properties of *mev-1* trend in the same direction as those of PB306. If the elevated endogenous oxidative stress experienced by *mev-1* is reflective of environmental stress in general, it suggests that the lab environment may be slightly stressful for PB306 compared to the lab-adapted N2. Moreover, the private allele spectrum of the wild isolates is closer to that of PB306 than of N2, which, by the preceding logic, suggests that the lab environment may be especially comfortable for N2 relative to the natural environment. The finding that there is too much variation in the standing mutation spectrum among wild isolates to be explained by a uniform mutational process suggests that the genetic variation in the mutational process observed between N2 and PB306 is not an artifact of the lab environment. Probably the most important point to take away from the comparison, however, is that both the rate and spectra of N2 and PB306 are not very different, which implies that conclusions about mutation drawn from N2 MA lines may be generalizable to *C. elegans* at large.

The third key finding is the discrepancy in the indel bias between the MA lines and the wild isolate private alleles: MA lines show a strong deletion bias (~62% genome-wide), whereas the deletion bias in the private alleles is much weaker (~54% deletion). As noted above, the indel rate has been shown to increase under stress in a variety of experimental systems, including *C. elegans*. The reduced deletion bias in *mev-1* relative to N2 reinforces the inference that a nematode’s life in nature is more stressful than life in the lab, with the attendant differences in the mutational process. To the extent that mutational variance in phenotypic traits is underlain by indel mutations, evolutionary inferences that rely on comparisons between mutational and standing genetic variance are rendered less robust.

The fourth key finding is the qualitative difference in the base-substitution mutation spectra between mononucleotide repeat sequence and non-mononucleotide sequence in the MA lines, and how the MA spectra relate to the standing private allele spectra. This study was predicated on the discrepancy between the (N2) MA base-substitution spectrum and that seen in wild isolates. In fact, when the non-mononucleotide spectra are compared, the discrepancy substantially disappears. Although significant differences remain, they do not appear wildly different **(Figure 4)**. Conversely, there is a huge discrepancy between the mononucleotide spectra of MA lines and private alleles, and in particular with respect to how A:T mutates. First, a larger fraction of mutations occur at A:T sites in the MA lines than in the private alleles (81% vs. 66%). Second, there are many more A:T→T:A transversions in the MA lines than in the private alleles (65% vs. 28%) and there are many fewer A:T→G:C transitions in the MA lines than in the private alleles (8% vs. 25%). The technology employed in sequencing most (not all) of the wild isolates and most (not all) of the MA lines was the same, and the data were analyzed in the same way (**Extended Methods, Supplemental Appendix A1.4; Supplemental Figure S4**), so it is difficult to imagine that the discrepancy is due to wildly different rates of false positives or false negatives in one or the other set of data. Given that the genetic basis of eukaryotic DNA repair is complex, and hundreds of genes are known to influence variation in DNA repair (Eisen and Hanawalt 1999), a reasonable guess is that some element(s) of DNA repair differs systematically between the lab environment and the natural environment, but what it may be we cannot say.

## Methods and Materials

### Strains

#### (i) MA lines

The *mev-1* gene encodes a subunit of succinate dehydrogenase cytochrome b, a component of complex II of the mitochondrial electron transport chain. The *mev-1(kn1)* allele is a G→A transition that replaces a glycine with a glutamate, resulting in decreased enzyme activity and increased electron leak. The resultant phenotype has been explored in several species (Ishii et al. 1990; Ishii et al. 2011; Ishii et al. 2013; Ishii et al. 2016) and includes elevated 8-oxodG and increased nuclear mutation rate (Hartman et al. 2004; Ishii et al. 2005). The *mev-1(kn1)* allele was backcrossed into the canonical Baer lab N2 strain genomic background; see **Extended Methods, Supplemental Appendix A1.1** for details.

PB306 is a wild isolate generously provided by Scott Baird; the N2 lines are derived from a replicate of the ancestor of the Vassilieva and Lynch (1999) MA lines.

#### (ii) Wild isolates

Genome sequence data for 773 unique wild isolates were obtained from the *C. elegans* Natural Diversity Resource (CeNDR; Cook et al. 2017). Details of the sequencing and variant calling are available at https://www.elegansvariation.org/.

### Mutation accumulation experiments

Details of the N2 and PB306 mutation accumulation experiment are given in Baer et al. (2005), those of the *mev-1* MA experiment are given in Joyner-Matos et al. (2011), and are summarized in the **Extended Methods, Supplemental Appendix A1.2**. Data on MA line transfers is presented in **Supplemental Table S1**. *mev-1* MA lines were propagated for up to 125 generations of MA (Gmax=125); N2 and PB306 lines were propagated for up to 250 generations of MA (Gmax=250). The basic protocol in both experiments follows that of Vassilieva and Lynch (1999) and is depicted in **Figure 1**.

### Genome sequencing and bioinformatics

32 N2 lines and 20 PB306 lines were previously sequenced with Illumina short-read sequencing at an average coverage of ~ 27X read-depth with 100 bp paired-end reads. The N2 data have been previously reported (Saxena et al. 2019). For this project we sequenced an additional 40 N2 lines, 54 PB306 lines, and 24 *mev-1* lines and their ancestors using Illumina short-read sequencing at an average coverage of 46X depth with 150 bp paired-end reads. Protocols for DNA extraction and construction of sequencing libraries are given in the **Extended Methods, Supplemental Appendix A1.3**; details of bioinformatics processing of raw sequence data and variant calling are given in the **Extended Methods, Supplemental Appendix A1.4**.

Following preliminary analysis, variants were called using HaplotypeCaller (in BP_RESOLUTION mode) in GATK4 (v4.1.4.0) (McKenna et al. 2010). Variants were identified as putative mutations if (1) the variant was identified as homozygous, and (2) it was present in one and only one MA line. Criterion (1) means that any mutations that occurred in the last few generations of MA that were still segregating and/or occurred during population expansion for DNA extraction were ignored. Because the MA progenitor was at mutation-drift equilibrium (Lynch and Hill 1986), the segregating variation is expected to be the same in the MA progenitor and the MA line, so ignoring heterozygotes results in an unbiased estimate of mutation rate.

Criterion (2) reduces the probability of mistakenly identifying a variant segregating at low frequency in the expanded population of the MA progenitor as a new mutation. One *mev-1* line was found not to carry the *mev-1(kn1)* allele and was omitted as a presumed contaminant. Four pairs of N2 strain MA lines and seven pairs of PB306 strain lines shared multiple variants and were inferred to have experienced contamination at some point during the MA phase; one line from each pair was arbitrarily omitted from subsequent analyses. The final data set includes 68 N2 lines, 67 PB306 lines, and 23 *mev-1* lines (**Supplemental Table S3**).

### Data Analysis

#### (i) Nuclear mutation rate

Because different lines experienced different numbers of generations during the MA phase, and because the number of callable sites differs among lines, we cannot use the number of mutations per line to directly compare mutation rates among strains. Instead, we calculated a mutation rate *μ_i_* for each line *i* as *μ_i_=m_i_/n_i_t_i_*, where *m* is the number of mutations, *n* is the number of callable sites, and *t* is the number of generations of MA (Denver et al. 2009). To test for group-specific effects on mutation rate, and to partition the (co)variance in mutation rate into within and among-group components, we consider the mutation rate itself as a continuously-distributed dependent variable in a general linear model (GLM). Details of the GLM analyses are given in the **Extended Methods, Supplemental Appendix A1.5.**

#### (ii) mtDNA mutation rate

Estimation of the mtDNA mutation rate is more complicated than for nuclear loci because a non-trivial fraction of mutations will not have reached fixation and remain heteroplasmic. The probability of a mutation ultimately reaching fixation is its current frequency in the population (Wright 1931), so the mtDNA mutation rate of MA line *i* can be estimated as 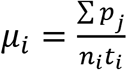, where *p_j_* is the frequency of the *j*’th mutation in line *i*, *n_i_* is the number of callable sites in line *i*, and *t_i_* is the number of generations of MA of line *i* (Konrad et al. 2017). However, since many lines have no mtDNA mutations, GLM is not an appropriate analysis. Instead, we used randomization tests of hypothesized differences in mtDNA mutation rate; the details are presented in the **Extended Methods, Supplemental Appendix A1.6**

#### (iii) Mutation spectrum

Mutation spectra were compared between groups by Fisher’s Exact Test. For comparisons with sample sizes too large to calculate directly we used Monte Carlo estimates of the FET as implemented in the FREQ procedure of SAS v.9.4.

We also tested the hypothesis that the variance in the private allele frequency spectrum among wild isolates can be explained by sampling variance around a single uniform base-substitution spectrum with expectation equal to the observed frequencies. We employed parametric bootstrap simulations, the details of which are presented in the **Extended Methods, Supplemental Appendix A1.7**.

## Supporting information

Supplemental Appendix A2. RIAILs methods

Supplemental Table S10_RIAIL genotypes

Supplemental Table S1_MA backup data

Supplemental Table S2_tests of fixed effects

Supplemental Table S3_Line-specific mutation rates

Supplemental Table S4_list of nuclear mutations with genomic context

Supplemental Table S5_mtDNA mutations

Supplemental Table S6_correlations of SNP and indel mutation rates

Supplemental Table S7_correlations of type-specific mutation rates

Supplemental Table S8_3bp motif-specific mutation rates

Supplemental Table S9_wild isolate variant counts

## Acknowledgments

We thank J. Dembek for laboratory assistance. Support was provided by NIH awards GM107227 to CFB and ECA, and GM127433 to CFB and V. Katju.

## Notes

### Competing Interest Statement

The authors have declared no competing interest.

### Summary of Updates

Supplemental Material added

