## Supplemental Appendix A1. Extended methods for "Mutability of mononucleotide repeats, not oxidative stress, explains the discrepancy between laboratory-accumulated mutations and the natural allele-frequency spectrum in *C. elegans*"

1. Strains. *(i) MA lines*. The *mev-1* gene encodes a subunit of succinate dehydrogenase cytochrome b, a component of complex II of the mitochondrial electron transport chain. The *mev-1(kn1)* mutation is a G→A transition that replaces a glycine with a glutamate, resulting in decreased enzyme activity and increased electron leak. The resultant phenotype has been explored in several species (Ishii et al. 1990; Ishii et al. 2011; Ishii et al. 2013; Ishii et al. 2016) and includes elevated 8-oxodG and mutation (Hartman et al. 2004; Ishii et al. 2005). The *mev-1(kn1)* allele was backcrossed into the canonical Baer lab N2 strain genomic background for 12 generations, followed by six generations of selfing by single-worm transfer to render the introgressed strain homozygous and to allow it to reach mutation-drift equilibrium. Consistent with previous reports (Ishii et al. 1990), the introgressed *mev-1* ancestor had lower total reproductive output and shorter lifespan relative to N2 (Joyner-Matos et al. 2011).

The sequence of the *mev-1* locus was inspected by eye in all *mev-1* MA lines. One line (735) did not have the *mev-1(kn1)* allele, presumably due to a cross-contamination event with an N2 strain, and was omitted from the analysis. All other *mev-1* lines were confirmed as homozygous for the *mev-1(kn1)* allele.

2. Mutation accumulation experiments**.**

Details of the N2 and PB306 mutation accumulation experiment are given in Baer et al. (2005), those of the *mev-1* MA experiment are given in Joyner-Matos et al. (2011). The basic protocol in both experiments follows that of Vassilieva and Lynch (1999) and is depicted in **Figure 1** in the main text. Briefly, replicate populations (MA lines) were initiated from a cryopreserved stock of a highly-inbred ancestor at mutation-drift equilibrium and maintained over the course of the experiment by transferring a single immature hermaphrodite at four-day intervals. Worms were maintained on 60mm NGM agar plates, spotted with 100 *μl* of an overnight culture of the OP50 strain of *E. coli* B, at a constant 20°C.

At every generation, the prior two generations of each MA line were kept at 20° C as backups; if the leading generation worm did not reproduce, the plate was reinitiated with an immature individual from the previous generation; this is referred to as "going to backup". If the parental worm had reproduced but no offspring had reached the L2 stage, we kept the offspring in the experiment to reproduce in the next generation; we refer to this delay in reproduction as "holding over". Holding over reduces the actual number of generations of evolution of an MA line below the maximum (Gmax). Ideally, going to backup should not affect the total number of generations of MA, because the backup worm should be a (double) first cousin of the worm that would have been transferred. However, there is some opportunity for overlapping generations on backup plates (e.g., if we had to go to backup twice or more in a row), so there is some ambiguity with respect to the true number of generations experienced by a line. The upper bound on the number of generations is the total number of possible transfers in the experiment (Gmax); the lower bound is the number of successful transfers (Gmin). Data on transfers and additional context is presented in **Supplemental Table S1**.

3. DNA Extraction and library preparation. Cryopreserved tubes of worms were thawed and grown for several days on 60 mm NGMA agar plates containing Streptomycin and Nystatin and seeded with 110 μl of overnight culture of the HB101 strain of *E. coli*. When food was nearly exhausted, a small piece (~50 mm^2^) of the agar plate was transferred onto a 100 mm NGMA plate and worms were allowed to grow for another 2-3 days. When food was nearly exhausted, worms were collected into 15 ml centrifuge tubes on ice and allowed to settle. Approximately 100 μl of settled worms were transferred into a 1.5 ml microcentrifuge tube and stored at -80°C until all samples were ready for DNA extraction, at which time samples were thawed and genomic DNA was extracted with the DNeasy Kit (Qiagen) following the manufacturer's protocol. Extracted DNA was stored at -20°C prior to library preparation.

Frozen DNA samples were thawed and diluted with diH_2_O to a concentration of 0.2 ng/μl. Tagmentation was performed per manufacturer’s instructions (Illumina, Nextera DNA Sample preparation kit, FC-121-1030). Following tagmentation, amplification was performed using custom barcoded IDT primers. Following amplification, a 96-pooled well sample was generated. Once pooled, 170µL of sample was combined with 30µL of 6X loading dye and run on a 2% agarose gel. DNA segments of 300-500bp were excised from gel. The QIAquick Gel Extraction Kit (QIAquick, 28704) was used to clean and elute DNA from the gel, following manufacturer’s instructions. Pooled library samples were quantified using the Qubit HS kit (Qubit, Q33230) and sequenced on a NovaSeq 6000 by Novogene, Inc.

4. Variant calling. (i) MA lines. Adapter sequence was trimmed from raw sequencing reads using fastp (Chen et al. 2018). Following trimming, we used bowtie2 (Langmead et al. 2009) to align trimmed sequence data to the N2 reference genome (WS263). Reads with the MQ<3 were removed from further analyses. Duplicate reads were identified and removed using MarkDuplicates tool in Genome Analysis Toolkit (GATK)/picard. Variants (SNPs and indels) were called using HaplotypeCaller (in BP_RESOLUTION mode) in GATK4 (v4.1.4.0) (McKenna et al. 2010). Next, the resulting files from all samples (23 *mev-1*, 68 N2, and 67 PB306 samples plus their ancestors) were consolidated using GenomicsDBImport, and variants called jointly using GenotypeGVCFs in GATK.

Variants that met the following three criteria were considered as putative mutations: (1) they were called homozygous; (2) they were present in one and only one MA lines for each datastore, and (3) the ancestor genotype is homozygous wild-type (Saxena al., 2019). We applied a 3x coverage threshold for filtering the sites with <3x coverage and include only sites that are covered >3X in > 90% of the MA lines in each strain (for example 62 out of 68 in N2 MA lines). Putative indels ≥ 20 bp and all indels within 50 bp of another putative indel were visually inspected using the Integrative Genomics Viewer (IGV) software.

Potential functional effects of mutations were assessed using snpEFF 4.3 (Cingolani et al. 2012), and the annotation database WBcel235.82. Each variant potentially has multiple effects; snpEFF sorts them by potential impact (i.e., the highest putative impact is listed first). We include only the largest potential effect of each variant (**Supplemental Table S3**).

(ii) Wild isolates. The details of the CeNDR variant calling pipeline are given on the CeNDR website (<https://www.elegansvariation.org/>). The only difference between the CeNDR variant-calling pipeline and our MA mutation-calling pipeline is that the CeNDR pipeline uses BWA (Li and Durbin 2010) as the mapping tool whereas our mutation-calling pipeline uses Bowtie2. To address the possibility that the discrepancy in the mutation spectra between MA lines and wild isolates can be explained by the use of different mapping software, we re-analyzed the *mev-1* MA data using the CeNDR pipeline (i.e., using BWA rather than Bowtie2 as the aligner). The resulting mutation spectra are nearly identical, both for base-substitutions and indels (**Supplemental Figure S4**). Thus, we conclude that the discrepancy between the MA spectra and the wild isolate spectra cannot be explained by the use of different mapping software.

5. Data Analysis: General Linear Models (GLM). Line-specific mutation rates were used as the dependent variable in analyses of mutation rate. The model can be formally written as *y_ijk_* = *μ + a_j_ + b_k_ +ab_ik_ + e_i|jk_*, where *y_ijk_* is the rate of mutation type *j* in line *i* of strain *k*, *μ* is the overall mean, *a_j_* is the fixed effect of mutation type *j*, *b_k_* is the fixed effect of strain *k*, *ab_bik_* is the fixed effect of the interaction between mutation type *j* and strain *k*, and *e_i|jk_* is the residual effect associated with MA line *i* and where it is assumed that *e* ~N(0,**R**), with covariance structure **R** (see below). Covariances were estimated by restricted maximum likelihood (REML); fixed effects were tested by F-tests with Type III sums of squares. Analyses were implemented using the MIXED procedure in SAS v. 9.4.

We compared models with different covariance structures (**R**) by finding the model that minimizes the corrected Akaike Information Criterion (AICc). If the best model by AICc includes more parameters than the next-best model, we compared the two by Likelihood Ratio Test (LRT). The models are nested, so twice the difference in the log-likelihoods is asymptotically chi-square distributed with degrees of freedom equal to the difference in the number of parameters estimated. We initially pooled the data across strains to find the best model, then compared that model to a model with the covariance structure estimated separately for each strain and tested by the same criteria. The covariance structures tested, in increasing order of complexity, are: variance components (a single, common variance and all covariances constrained to equal zero), compound symmetry (common variance, common covariance); banded main diagonal (a separate variance estimated for each variable, all covariances constrained to zero), and unstructured (all elements of the covariance matrix estimated individually).

Preliminary analyses revealed qualitatively different mutation rates and spectra for mononucleotide repeats (defined as five or more consecutive bases of the same type) compared to all other sequence motifs; see section 8 below for details of the bioinformatics by which mononucleotide repeats were identified. Rather than extend the above GLM to include sequence type (mononucleotide and non-mononucleotide), which would have included three two-way interactions as well as the three-way interaction, we broke the analysis down into two different two-factor models. First, we analyzed each of the six base-substitution rates individually, including strain and sequence type as predictor variables. Comparisons among strains were restricted to the comparison between *mev-1* and N2 (the foundational motivation for the study) and the comparison between N2 and PB306. Next, since in only one case did the strain-by-sequence type interaction approach statistical significance (see Results), we pooled the data over strains (N2 and PB306, omitting *mev-1*) and analyzed the model with sequence type and mutation type as the predictor variables.

6. mtDNA mutation rate. Estimation of the mtDNA mutation rate is more complicated than for nuclear loci because a non-trivial fraction of mutations will not have reached fixation and remain heteroplasmic. The probability of a mutation ultimately reaching fixation is its current frequency in the population (Wright 1931), so the mtDNA mutation rate of MA line *i* can be estimated as

$\mu_{i}=\frac{{\sum p}_{j}}{n_{i}t_{i}}$, where *p_j_* is the frequency of the *j*'th mutation in line *i*, *n_i_* is the number of callable sites in line *i*, and *t_i_* is the number of generations of MA of line *i* (Konrad et al. 2017).

Because many lines have no mtDNA mutations, the residuals of mtDNA mutation rate are far from normally distributed, so GLM is not an appropriate method by which to test the hypothesis that mtDNA mutation rates differ among strains. Instead, we used a randomization test, in which (1) the number of lines/strain is held constant and the data (line-specific mutation rates) randomly shuffled among strains, (2) the average mutation rate calculated for each strain, and (3) the difference in average mutation rate between the three pairs of strains calculated. This procedure was repeated 1000 times, to generate a null distribution of pairwise differences in mutation rate. If the observed value of the pairwise difference is greater than 95% of the randomized values, we take it to be statistically significant at P<0.05.

7. Parametric bootstrap simulations of wild isolate mutation spectra. To test the hypothesis that the variance in the private allele frequency spectrum among wild isolates can be explained by sampling variance around a single uniform base-substitution spectrum with expectation equal to the observed frequencies, we employed parametric bootstrap simulations, as follows. First, each wild isolate was assigned its observed number of private alleles, sampled with replacement with probability equal to the observed frequency (e.g., if the frequency of A:T→T:A base substitutions is 20%, the probability that any simulated private allele is the result of a A:T→T:A mutation is 20%). Next, the Kullback-Leibler divergence (Kullback and Leibler 1951), $D_{KL}=\sum_{i=1}^{n} p_{i}log\frac{p_{i}}{q_{i}}$, was calculated for each wild isolate, where *p_i_* and *q_i_* are the observed and expected base-substitution frequencies, and the mean divergence $\bar{D}_{KL}$ of the set of wild isolates calculated, as well as the variance in *D_KL_* among wild isolates. This procedure was repeated 1000 times to generate a simulated frequency distribution of the mean *D_KL_* and its variance under the null hypothesis, and the observed values compared to the null distribution. If the observed value is greater than 95% of simulated values, we reject the null hypothesis at the p<0.05 level.

8. Mononucleotide repeats. We used Phobos 3.3.12 software (http://www.rub.de/ecoevo/cm/cm_phobos.htm) to identify all mononucleotide repeats of ≥ 5 bp in the WS263 build of the *C. elegans* genome. To identify imperfect repeats, we applied the default mismatch penalties (mismatch score -5, gap score -5, recursion depth 5, Minimum score = 0). The minimum repeat unit criterion was 5bp and we kept all the sites with 100% repeat perfection. We increased the range of all mononucleotide repeat by two bases on either end of the repeat, following Saxena et al. (2019). The average coverage of mononucleotide repeats and non-mononucleotide repeat in the MA lines of the three strains was estimated using mosdepth 0.3.1 (Pedersen and Quinlan 2018); the same coverage threshold was applied for mononucleotide and non-mononucleotide regions. Including the 2 bp flanking regions, mononucleotide repeats include 14,470,735 nucleotides and non-mononucleotide regions include 85,815,671 nucleotides. Excluding the flanking regions, mononucleotide repeats *per se* include 8,578,075 nucleotides. The GC-content of mononucleotide and non-mononucleotide sequence is 20.12% and 38.02%, respectively.

9. Local sequence context. We counted the total number of each of the 64 3-base motifs in the reference genome using Jellyfish v. 2.3.0 software (Marcais and Kingsford 2011). For example, in the sequence -5'-AAAA-3', there are two possible 5'-aaA-3' motifs. We applied a custom Bash script to count the number of each motif in the MA lines. We used bcftools to extract the position of each variant, and bedtools to extract the sequence context of each variant.

We counted the total number of mutations occurring at the 3' position in each of the 3-bp motifs and estimated the base-substitution mutation rate of each 3-base motif *i* in an MA line as $\mu_{i}=\frac{X_{i}}{n_{i}t}$, where *X* is the total number of mutations in each 3-bp motif in the MA line, *n* is the detectable number of each 3-base motif in that line, and *t* is the number of generations of MA. The reported values of *μ_i_* are unweighted means over all lines in the strain. Scripts are deposited in github at https://github.com/moeinrajaei/mev-1-project.

10. False positives (FP) and false negatives (FN).

*(i) False Positives*. False positives were assessed in two ways. First, we counted sites that differ between our MA ancestor and the reference (N2) genome (i.e., sites that are scored 1/1 in the MA ancestor and 0/0 in the reference genome). These are presumed to be new mutations fixed subsequent to the divergence of the MA ancestor and the reference strain from their common ancestor. These sites were scored in each MA line in the set. If a site is scored 1/1 in the MA ancestor but scored 0/0 in all MA lines, we inferred a false positive in the MA ancestor. The power of this method is obviously sensitive to the number of MA lines sequenced; the more MA lines sequenced, the greater the chance of identifying a false positive. Since the number of N2 lines (68) and PB306 lines (67) are nearly equal, any difference in the frequency of false positives cannot be attributed to differences in power.

Second, we employed an independent set of low-coverage sequence data from a set of 192 recombinant inbred advanced intercross lines (RIAILs) generated from a cross between two N2 strain MA lines, 530 and 563. The details of the construction and sequencing of the RIAILs and data analysis are presented in **Supplemental Appendix A2**.

*(ii) False negatives*. To identify false negatives, we first introduced simulated variants ("dummy mutations") into the reference genome, and then re-analyzed the MA data using the simulated reference genome. For example, suppose position *x* in the reference genome is A. We randomly change the A to a C, i.e. an A →C "dummy" transversion. In this case, the MA ancestor and all MA lines should have a C at site *x*. If the genotype at site *x* in any MA line is not called a C/C homozygote, for any reason, it is classified as a failure to recall. If the genotype at site *x* in an MA line was called "not C/C" (i.e., it was scored A/A or A/C), it is a false negative. We introduced 983 dummy base substitutions into the reference genome; failure to recall rates are given in the main text.

Indels present a more complex problem, because inserting or deleting sequence changes the physical map of the reference genome relative to the focal genome. To address the question of indel false negatives, we first introduced 9,990 dummy indels into the reference genome (**Supplemental Table S11** below). Next, we defined five size bins of 1-5 bp, 6-10 bp, 11-20bp, 21-50bp, and >50 bp, and analyzed the data as described above for base-substitutions. For each MA line, we counted the total number of indels in each bin using the simulated reference genome and the true reference genome and compared the number of indels in each bin. We then repeated the simulation and analyses with a second set of dummy mutations; the correlation between the line-specific failure to recall rates in the two simulations provides an estimate of the reliability of the method (**Supplemental Figure S5** below).

| **Bin (bp)** | **Deletions** | **Insertions** | **Total Indels** |
| --- | --- | --- | --- |
| 1-5 | 668 | 707 | 1375 |
| 6-10 | 660 | 680 | 1340 |
| 11-20 | 1034 | 1130 | 2164 |
| 21-50 | 1898 | 1922 | 3820 |
| >50bp | 639 | 652 | 1291 |

**Table S11**. Size distribution of the dummy indels introduced into the reference genome.

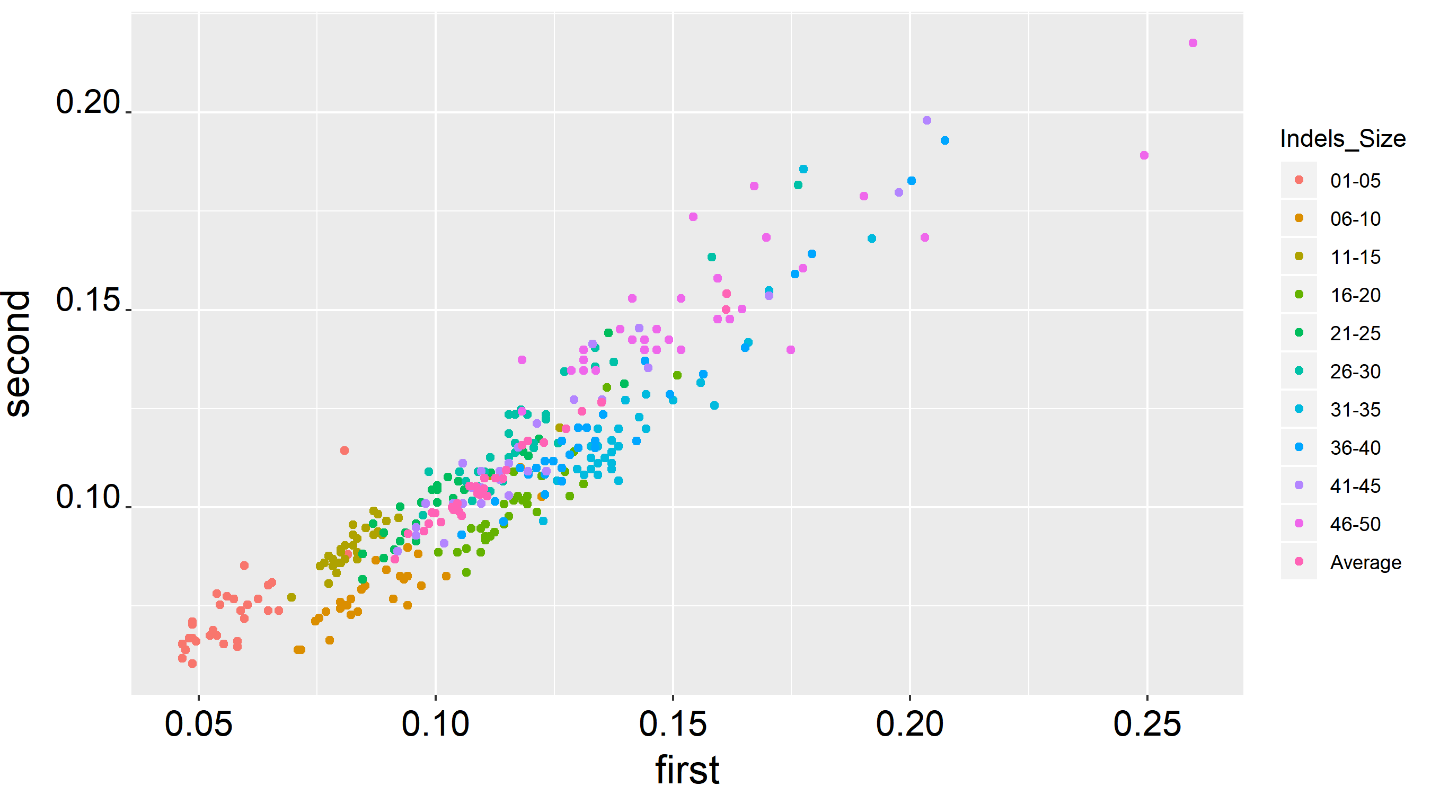

**Figure S5.** Plot of the failure to recall rates of the two simulated dummy data sets; the first simulation is on the x-axis, the second simulation is on the y-axis (r=0.93, P << 0.0001). Each data point of a given color represents an individual MA line (n=30 simulated lines).

11. Indel filters

To compare the indel spectra of MA lines and segregating private alleles, we applied the following filtering criteria using VariantFiltration in GATK (4.1.4.0). Variants with quality (QUAL) < 30, quality by depth (QD) < 20, read depth (DP) < 5, strand odds ratio (SOR) > 5, Fisherstrand (FS) > 100, ReadPosRankSum value < -20. Sites with > 10% missing genotypes and or any heterozygous genotypes were removed.
