## Supplemental Appendix A2. RIAILs methods for "Mutability of mononucleotide repeats, not oxidative stress, explains the discrepancy between laboratory-accumulated mutations and the natural allele-frequency spectrum in *C. elegans*"

**Appendix A2**. Construction and analysis of recombinant inbred advanced intercross lines (RIAILs).

1. Line Construction. Two N2-strain MA lines, line 530 (243 MA generations) and line 563 (237 MA generations) were chosen as parents to generate a panel of 200 recombinant inbred advanced intercross lines (RIAILs) (Rockman and Kruglyak 2008). In brief, we used a random pair mating design with equal contributions of each parent to the next generation over ten generations of intercrossing, followed by ten generations of selfing. We first thawed the cryopreserved parental strains, lines 530 and 563, on 100mm plates seeded with OP50 *E. coli*. Both strains contained males when thawed, and we allowed them to propagate until many gravid hermaphrodites were present. We then bleached the strains and incubated the recovered embryos in M9 buffer overnight to make synchronized cultures of arrested L1 stage larvae. We plated these L1s on multiple 100 mm plates to avoid overcrowding and grew them for 48 hours. We then set up 30 independent outcrosses on mating plates (30mm plates seeded with 10uL of OP50 *E. coli*) with the sex of the parental strains reversed for half the outcrosses. We set up these and all subsequent crosses with two young males and one virgin hermaphrodite each, and always transferred the parents to a new mating plate two days after setting up the cross to avoid accidentally selecting self progeny. We made 280 independent crosses among F1 progeny, being careful to select animals that would provide equal proportions of the founders’ cytoplasmic and X chromosomes to the F2 progeny. Of these crosses, only 200 showed the roughly equal sex ratios in the progeny indicative of successful mating, therefore all subsequent crosses were made in duplicate to avoid failed matings in subsequent generations. We randomly paired F2 progeny from the 200 successful F1 crosses to generate the F3 generation and performed each cross reciprocally to maintain a constant population size of 200. We repeated this process over ten generations of intercrossing. In some cases, both duplicate crosses failed, in which case we randomly selected another line of the same generation to take its place, leading to the occasional overrepresentation of some parents within a generation of intercrossing. At the F10 generation we chose a single hermaphrodite from each of the 200 recombinant lines and allowed them to self fertilize for 10 generations. Of the 200 RIAILs, 196 survived cryopreservation and were used in this study.

2. Illumina library construction and whole-genome sequencing. To isolate genomic DNA from the recombinants, we transferred nematodes from two freshly starved 10 cm NGM plates into a 15 ml conical tube by washing with 10 mL of M9 buffer. We allowed the worms to settle by gravity the bottom of the conical tube, removed the supernatant, and added 10 mL of fresh M9. We repeated this wash method three times over the course of one hour to serially dilute any remaining *E. coli* in the M9 and allow the nematodes time to purge ingested bacteria. We then isolated genomic DNA from 100 to 300 µl nematode pellets using the Blood and Tissue DNA isolation kit cat# 69506 (QIAGEN, Valencia, CA) following established protocols (Cook et al. 2016). We quantified the concentration of DNA in each sample with the Qubit dsDNA Broad Range Assay Kit cat# Q32850 (Invitrogen, Carlsbad, CA) and diluted all samples to 0.2 ng/μL. We then incubated each sample with diluted Illumina transposome cat# FC-121-1031 (Illumina, San Diego, CA) and amplified the tagmented fragments with barcoded primers. We combined the uniquely indexed samples into 96-sample pooled libraries and size-selected them by separating the material on a 2% agarose gel, extracting the fragments ranging from 400-600 bp, and purifying the extraction with the QIAquick Gel Extraction Kit cat# 28706 (QIAGEN, Valencia, CA). We determined the concentration and size distribution of the purified, size-selected libraries with the 2100 Bioanalyzer (Agilent, Santa Clara, CA) before submitting them to the NUSeq Core Facility of Northwestern University where they were sequenced on the Illumina HiSeq4000 platform (paired-end 150 bp reads).

3. Variant calling and analysis. Variants were called using the same analysis pipeline as for the MA lines, described in the **Extended Methods, Supplemental Appendix A1.4**. We identified 699 SNPs and 1054 indels in the two parental lines (Note: these variants include new mutations in the MA phase and fixed mutations that occurred on the lineage leading to the MA ancestor subsequent to its divergence from its common ancestor with the reference genome). The variants were recalled jointly in 192 RIAILs and 636 SNPs and 961 indels were kept.

The expected frequency of a variant in the set of RIAILs is 50% (i.e., we expect Mendelian segregation). If a variant is not present in at least ten (out of 192) RIAILs and there are fewer than 50 RIAILs with missing data at the site, it is considered a false positive.
