## Supplemental Figures for "Mutability of mononucleotide repeats, not oxidative stress, explains the discrepancy between laboratory-accumulated mutations and the natural allele-frequency spectrum in *C. elegans*"

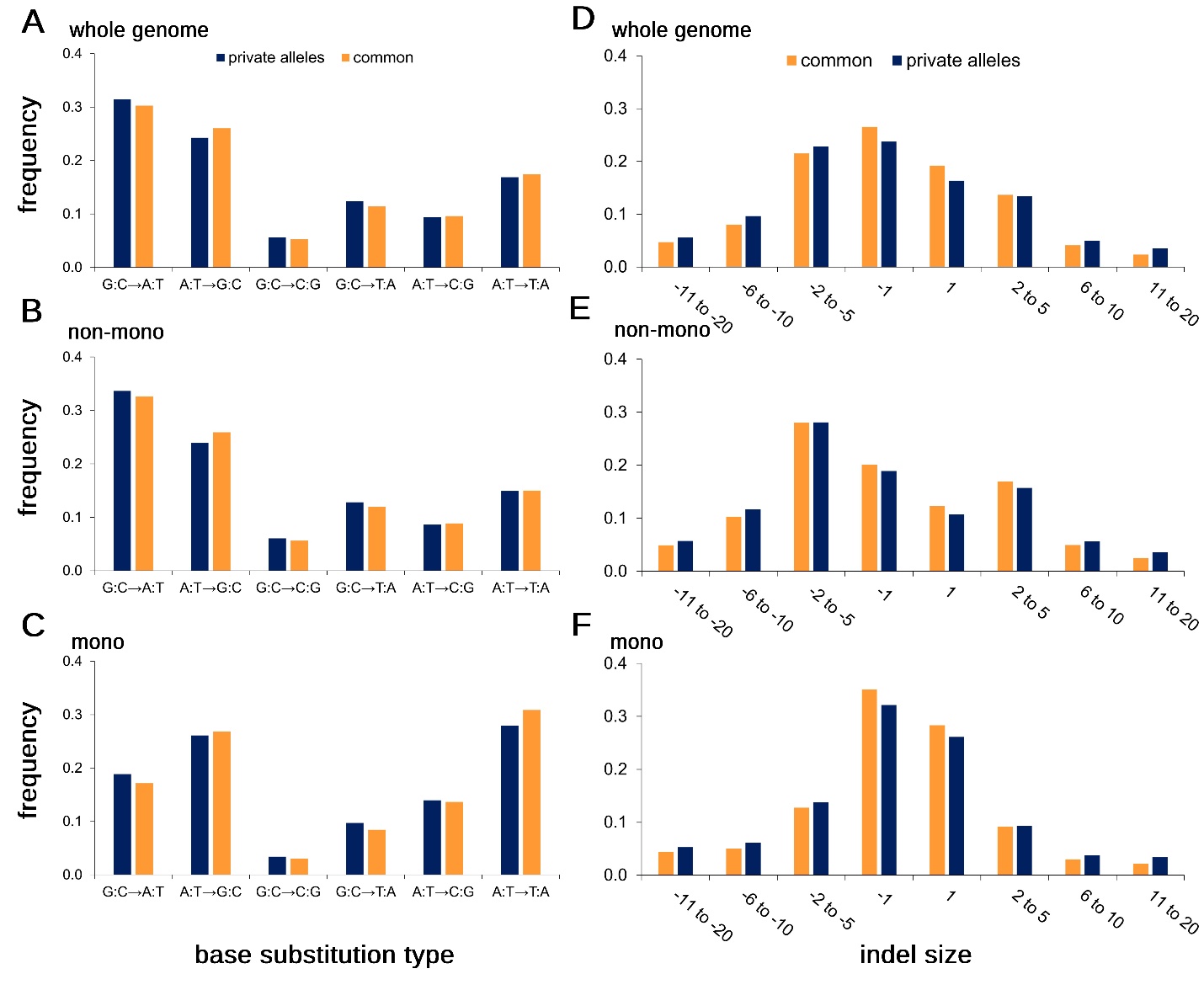

**Figure S1. Spectra of private alleles (n=1/N, blue) and common variants (orange)**. Left (**A-C**), Base-substitution spectra. Right (**D-F**), Indel spectra. Top panels (**A,D**), whole-genome; middle panels (**B,E**), non-mononucleotide sequence; bottom panels (**C,F**), mononucleotide sequence.

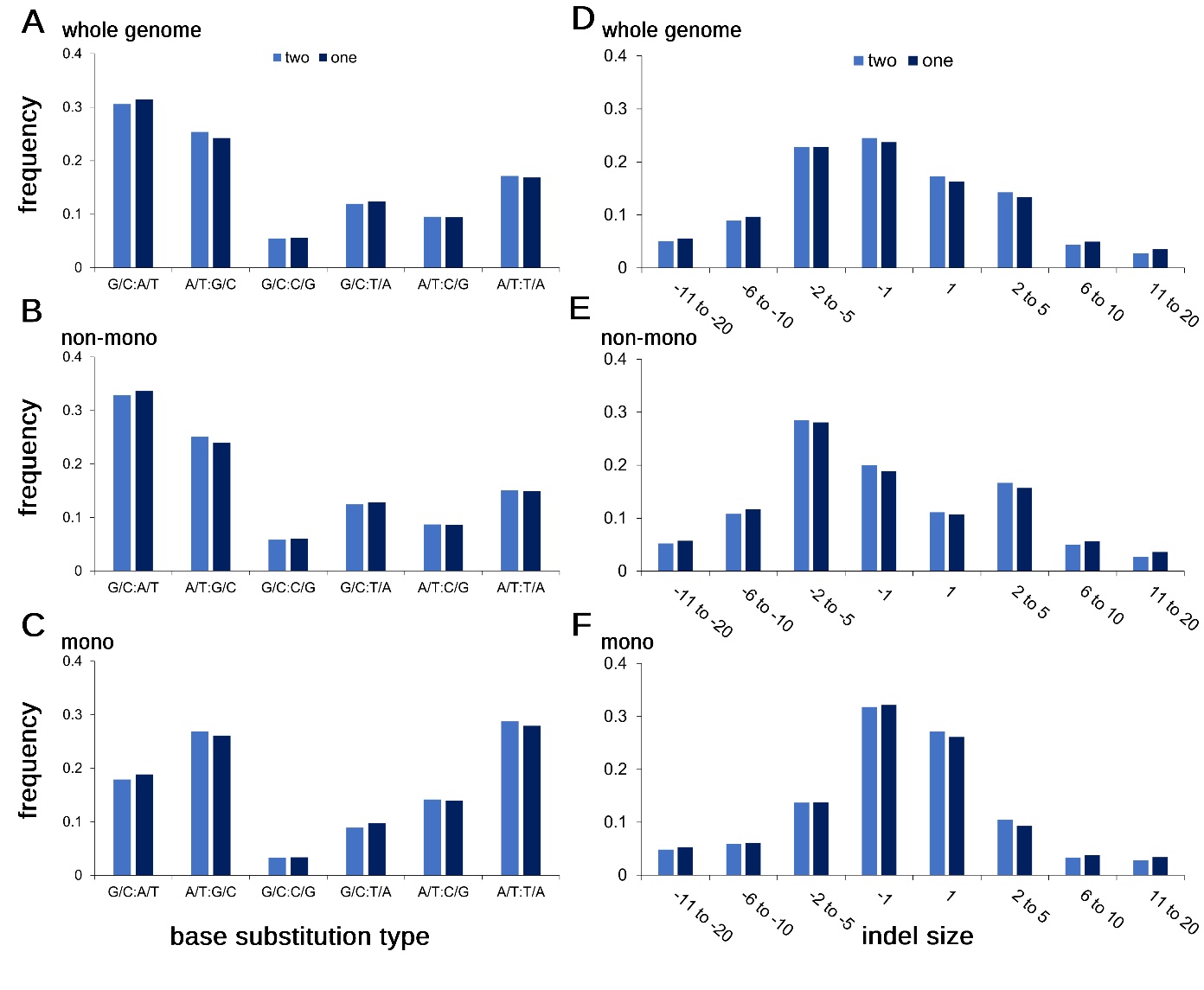

**Figure S2. Spectra of segregating singleton (dark blue, n=1/N) and doubleton (light blue, n=2/N) variants.** Left (**A-C**), base-substitution spectra. Right (**D-F**), indel spectra. Top panels (**A,D**), whole-genome; middle panels (**B,E**), non-mononucleotide sequence; bottom panels (**C,F**), mononucleotide sequence.

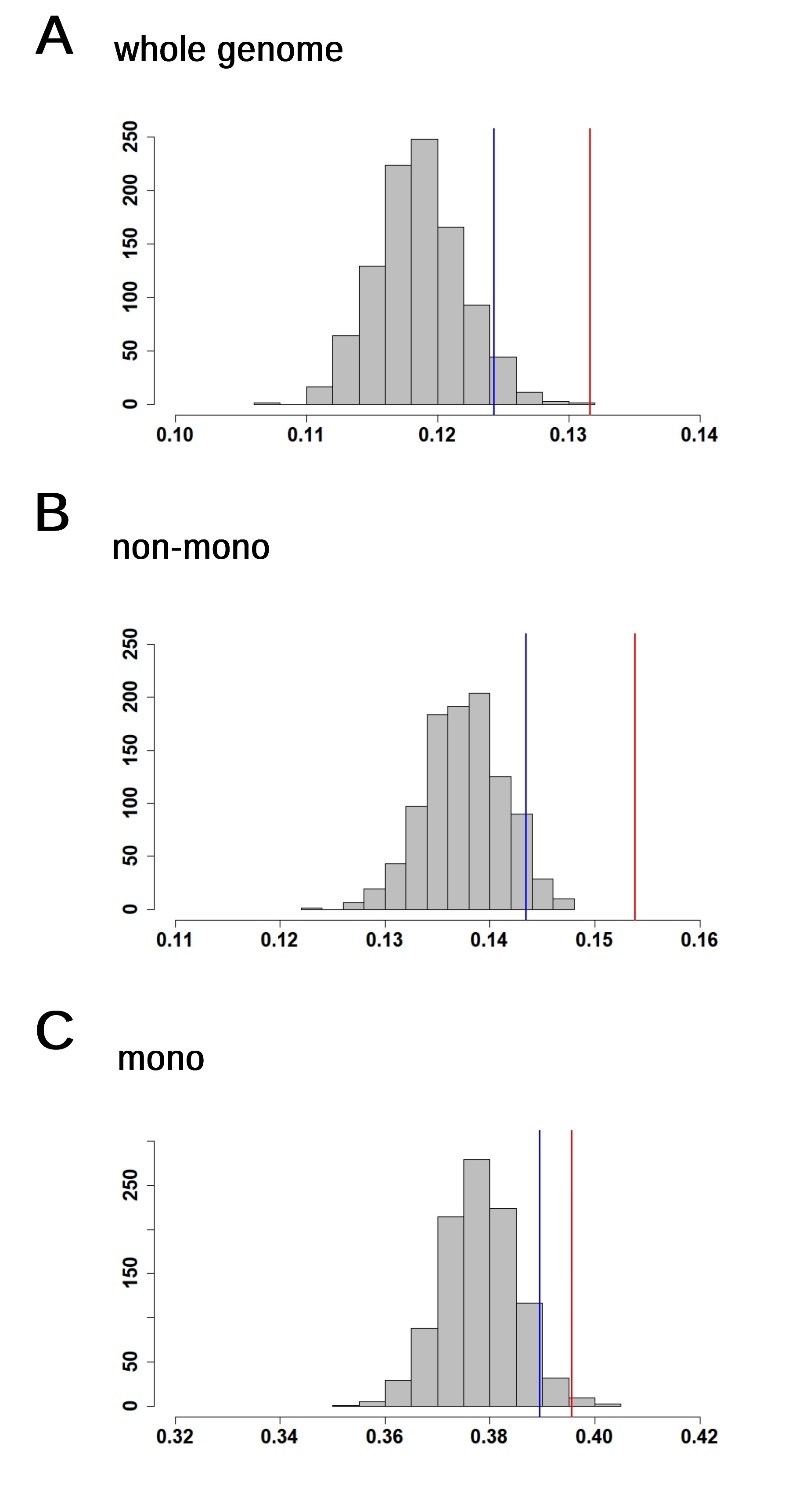

**Figure S3. Parametric bootstrap distributions.** Red lines show the observed values, blue lines show the upper 95% confidence limit of simulated D(KL).
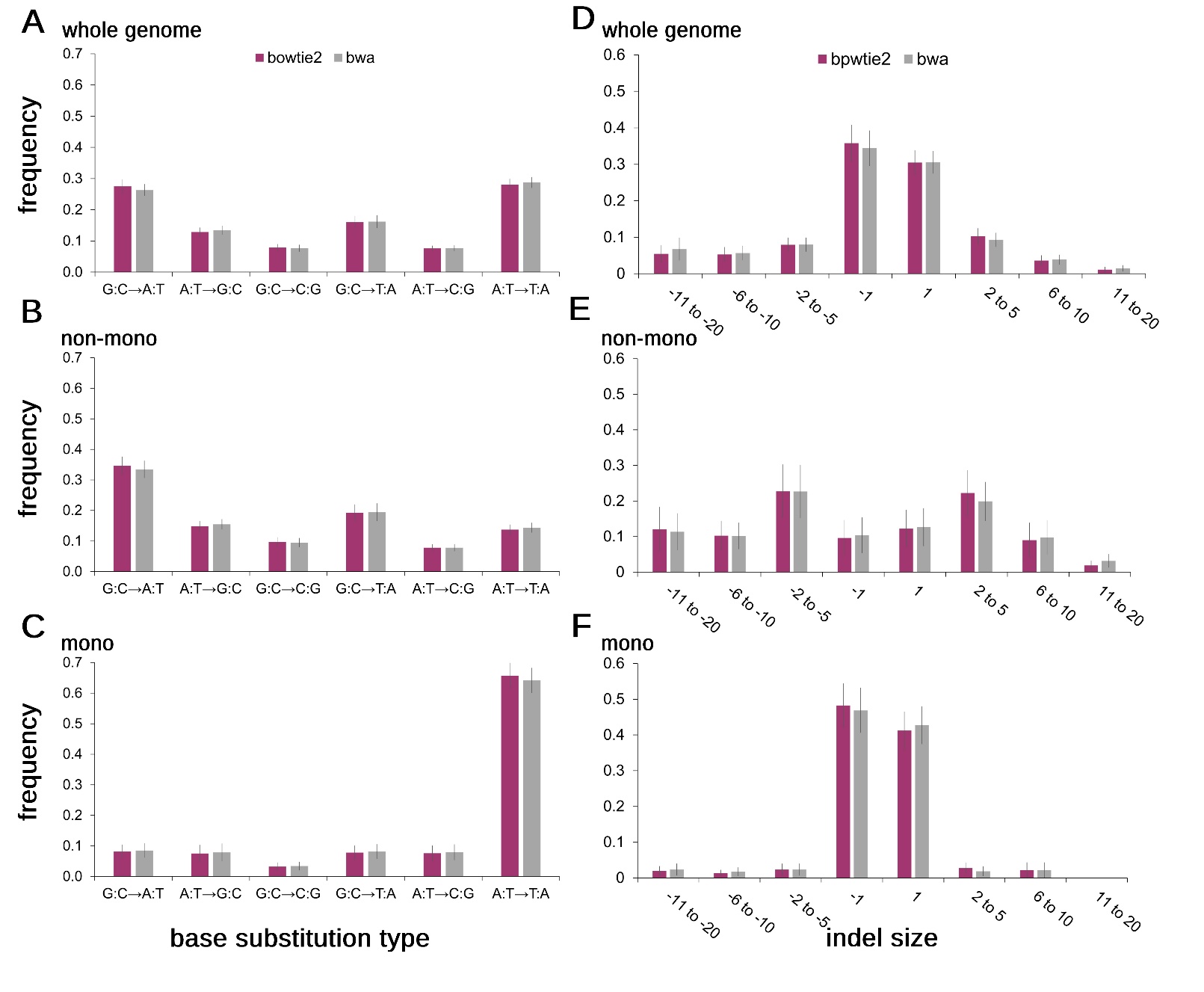
**Figure S4. Base-substitution and indel spectra of *mev-1* MA lines using two different mapping programs, Bowtie2 (maroon) and BWA (gray).** Left (**A-C**), base-substitution spectra. Right (**D-F**), indel spectra. Top panels (**A, D**) whole-genome; middle panels (**B, E**) non-mononucleotide sequence; bottom panels (**C, F**) mononucleotide sequence. Error bars show 1 SEM.
