## Supplemental Table S2_tests of fixed effects for "Mutability of mononucleotide repeats, not oxidative stress, explains the discrepancy between laboratory-accumulated mutations and the natural allele-frequency spectrum in *C. elegans*"

**Supplemental Table S2. Tests of fixed effects.** See Appendix A1.5 for details of the general linear model (GLM) and Table 1 in the main text for trait values. The columns in tables below are: 1. Description of the fixed effect; 2. Degrees of freedom, determined by the Kenward-Roger method; 3. F-statistic; 4. P-value, not corrected for multiple tests. Further descriptions are underneath the tables.

| **A**. Fixed Effect (*μ_BS_*, by type) | DF (num, den) | F | Pr>F |
| --- | --- | --- | --- |
| Base-substitution type (6) | 5,130 | 267.1 | <0.0001 |
| Strain (3) | 2,222 | 5.25 | 0.0059 |
| Strain x base-sub type | 10,198 | 2.48 | 0.0081 |

**A**. Genome-wide base substitution mutation rates, *μ_BS_*, including all three strains of MA lines, the six types of base-substitution, and the interaction.

| **B**. Fixed Effect (*μ_BS_*, pooled) | DF (num, den) | F | Pr>F |
| --- | --- | --- | --- |
| Strain (*mev-1* vs N2) | 1,56.8 | 15.4 | 0.0002 |

**B**. Pairwise-test of pooled genome-wide base-substitution rate difference between *mev-1* and N2.

| **C.** Fixed Effect (*μ_BS_*, pooled) | DF (num, den) | F | Pr>F |
| --- | --- | --- | --- |
| Strain (*mev-1* vs PB306) | 1,49.1 | 7.88 | 0.0072 |

**C.** Pairwise-test of pooled genome-wide base-substitution rate difference between *mev-1* and PB306.

| **D.** Fixed Effect (*μ_BS_*, pooled) | DF (num, den) | F | Pr>F |
| --- | --- | --- | --- |
| Strain (N2 vs PB306) | 1,131 | 2.10 | 0.1494 |

**D.** Pairwise-test of pooled genome-wide base-substitution rate difference between N2 and PB306.

| **E.** Fixed Effect (*μ_GC→TA_*) | DF (num, den) | F | Pr>F |
| --- | --- | --- | --- |
| Strain (all 3) | 2,55.1 | 4.09 | 0.0221 |

**E.** Planned comparison among strains of the GC→TA transversion rate.

| **F.** Fixed Effect (*μ_AT→TA_*) | DF (num, den) | F | Pr>F |
| --- | --- | --- | --- |
| Strain (all 3) | 2,58.7 | 4.25 | 0.0189 |

**F.** Post hoc comparison among strains of AT→TA transversion rate.

| Fixed Effect (*μ_INS_*) | DF (num, den) | F | Pr>F |
| --- | --- | --- | --- |
| **G.** Strain (all 3) | 2, 53.9 | 5.62 | 0.0061 |

**G.** Comparison among strains of the genome-wide insertion rate.

| **H.** Fixed Effect (*μ_DEL_*) | DF (num, den) | F | Pr>F |
| --- | --- | --- | --- |
| Strain (all 3) | 2,60.8 | 5.74 | 0.0052 |

**H.** Comparison among strains of the genome-wide deletion rate.

| **I.** Fixed Effect (*μ_INS_*) | DF (num, den) | F | Pr>F |
| --- | --- | --- | --- |
| Strain (*mev-1* vs N2) | 1, 26.9 | 8.06 | 0.0085 |

**I.** Pairwise test of genome-wide insertion rate difference between *mev-1* and N2.

| **J.** Fixed Effect (*μ_DEL_*) | DF (num, den) | F | Pr>F |
| --- | --- | --- | --- |
| Strain (*mev-1* vs N2) | 1, 34.4 | 0.26 | 0.6129 |

**J.** Pairwise test of genome-wide deletion rate difference between *mev-1* and N2.

| **K.** Fixed Effect (*μ_INS_*) | DF (num, den) | F | Pr>F |
| --- | --- | --- | --- |
| Strain (*mev-1* vs PB306) | 1, 31.2 | 2.42 | 0.1295 |

**K.** Pairwise test of genome-wide insertion rate difference between *mev-1* and PB306.

| **L.** Fixed Effect (*μ_DEL_*) | DF (num, den) | F | Pr>F |
| --- | --- | --- | --- |
| Strain (*mev-1* vs PB306) | 1, 50.1 | 4.13 | 0.0474 |

**L.** Pairwise test of genome-wide deletion rate difference between *mev-1* and PB306.

| **M.** Fixed Effect (*μ_INS_*) | DF (num, den) | F | Pr>F |
| --- | --- | --- | --- |
| Strain (N2 vs PB306) | 1, 122 | 5.37 | 0.0221 |

**M.** Pairwise test of genome-wide insertion rate difference between N2 and PB306.

| **N.** Fixed Effect (*μ_DEL_*) | DF (num, den) | F | Pr>F |
| --- | --- | --- | --- |
| Strain (N2 vs PB306) | 1, 115 | 11.48 | 0.0010 |

**N.** Pairwise test of genome-wide insertion rate difference between N2 and PB306.

| **O.** Fixed Effect (*μ_BS_*) | DF (num, den) | F | Pr>F |
| --- | --- | --- | --- |
| Strain (all 3) | 2, 272 | 0.89 | 0.4132 |
| **sequence type (non-mono vs. mono)** | **1, 243** | **120.92** | **<0.0001** |
| Strain x seq. type | 2, 272 | 0.15 | 0.8611 |
| base-sub type (all 6) | 5, 140 | 146.34 | <0.0001 |
| Strain x base-sub type | 10, 228 | 1.95 | 0.0399 |
| **seq. type x base-sub type** | **5, 140** | **86.49** | **<0.0001** |
| Strain x seq. type x base-sub type | 10, 228 | 1.16 | 0.3202 |

**O.** Test of effects of sequence type (non-mononucleotide vs. mononucleotide), strain, base-substitution type, and all interactions on the base-substitution rate (*μ_BS_*). The important comparisons are the ones highlighted in bold font.

| **P.** Fixed Effect (*μ_AT→TA_*) | DF (num, den) | F | Pr>F |
| --- | --- | --- | --- |
| Strain (all 3) | 2, 61.3 | 4.83 | 0.0113 |
| **sequence type (non-mono vs. mono)** | **1, 51.6** | **464.99** | **<0.0001** |
| Strain x seq. type | 2, 61.3 | 3.99 | 0.0235 |

**P.** Post-hoc test of the effect of sequence type (non-mononucleotide vs. mononucleotide), strain, and their interaction on the rate of AT→TA transversions.

| **Q.** Fixed Effect (*μ_1BP_INDEL_*) | DF (num, den) | F | Pr>F |
| --- | --- | --- | --- |
| Strain (all 3) | 2, 105 | 2.15 | 0.1217 |
| **sequence type (non-mono vs. mono)** | **1, 78.3** | **490.15** | **<0.0001** |
| Strain x seq. type | 2, 105 | 1.94 | 0.1482 |
| Indel type (+/- 1 bp) | 1, 78.3 | 8.68 | 0.0042 |
| Strain x indel type | 2, 105 | 0.66 | 0.5172 |
| seq. type x indel type | 1, 78.3 | 7.02 | 0.0097 |
| Strain x seq. type x indel type | 2, 105 | 0.60 | 0.5525 |

**Q.** Test of the effect of sequence type (non-mononucleotide vs. mononucleotide, strain, indel type (insertion vs. deletion), and their interactions on the rate of +/- 1 bp indels.

| **R.** Fixed Effect (*5'-ttA-3'*) | DF (num, den) | F | Pr>F |
| --- | --- | --- | --- |
| Strain (all 3) | 2, 57.6 | 4.78 | 0.0120 |
| **sequence type (non-mono vs. mono)** | **1, 48.5** | **210.49** | **<0.0001** |
| Strain x seq. type | 2, 57.6 | 4.43 | 0.0162 |

**R.** Test of the effect of sequence type (non-mono vs. mono), strain, and their interaction on the rate of 5'-ttA-3' mutation.
