## Supplemental Table S6_correlations of SNP and indel mutation rates for "Mutability of mononucleotide repeats, not oxidative stress, explains the discrepancy between laboratory-accumulated mutations and the natural allele-frequency spectrum in *C. elegans*"

x

| (**a**) ***mev-1*** | Del | Ins | SNP |
| --- | --- | --- | --- |
| Del |  | -0.21 | 0.21 |
| Ins |  |  | -0.025 |
| (**b**) **N2** | Del | Ins | SNP |
| Del |  | -0.065 | 0.24 |
| Ins |  |  | -0.07 |
| (**c**) **PB306** | Del | Ins | SNP |
| Del |  | 0.35 | 0.32 |
| Ins |  |  | 0.09 |

**Supplemental Table S6**. Correlations between base-substitution (SNP) and indel mutation rates. (a) *mev-1*; (b) N2; (c) PB306. Correlations are reported for each strain separately because the best-fit linear model includes the among-line covariance estimated separately for each strain. See Methods for details.
