## Supplemental Table S7_correlations of type-specific mutation rates for "Mutability of mononucleotide repeats, not oxidative stress, explains the discrepancy between laboratory-accumulated mutations and the natural allele-frequency spectrum in *C. elegans*"

x

| **Mut type** | **AT>GC** | **AT>TA** | **GC>AT** | **GC>CG** | **GC>TA** | **Row Ave** |
| --- | --- | --- | --- | --- | --- | --- |
| **AT>CG** | 0.27 | 0.10 | 0.19 | -0.03 | -0.06 | 0.09 |
| **AT>GC** |  | 0.26 | 0.20 | 0.07 | -0.14 | 0.13 |
| **AT>TA** |  |  | 0.17 | 0.06 | -0.03 | 0.11 |
| **GC>AT** |  |  |  | 0.00 | -0.01 | 0.11 |
| **GC>CG** |  |  |  |  | 0.09 | 0.04 |
| **GC>TA** |  |  |  |  |  | -0.03 |

**Supplemental Table S5**. Correlation matrix of the six type-specific base-substitution mutation rates. Data are pooled across strains because the best-fit linear model includes a single (pooled) estimate of the among-line covariance. See Methods for details. Two outliers were removed prior to analysis.
